# Integrative single-cell transcriptomics clarifies adult neurogenesis and macroglia evolution

**DOI:** 10.1101/2023.02.27.530203

**Authors:** David Morizet, Isabelle Foucher, Alessandro Alunni, Laure Bally-Cuif

## Abstract

Macroglia fulfill essential functions in the adult vertebrate brain, producing and maintaining neurons and regulating neuronal communication. However, we still know little about their emergence and diversification. We used the zebrafish *D. rerio* as a distant vertebrate model with moderate glial diversity as anchor to reanalyze datasets covering over 600 million years of evolution. We identify core features of adult neurogenesis and innovations in the mammalian lineage with a potential link to the rarity of radial glia-like cells in adult humans. Our results also suggest that functions associated with astrocytes originated in a multifunctional cell type fulfilling both neural stem cell and astrocytic functions before these diverged. Finally, we identify conserved elements of macroglial cell identity and function and their time of emergence during evolution.

**One-Sentence Summary:** Radial glia of the adult zebrafish forebrain associate transcriptomic features of adult neural stem cells and astrocytes

## Introduction

The appearance of nervous systems during animal evolution was a transformative event which revolutionized their interactions with the outside world and ultimately gave rise to higher cognition. To acquire their current forms, nervous systems required emergence of new cell types followed by substantial diversification(*1*). To date, work on cell type evolution in the brain has focused on neurons. Conversely, macroglia have been mostly overlooked, despite being a major component of the human nervous system, outnumbering neurons in the cerebral cortex(*2*). Macroglia fulfill essential roles to produce, guide and support neurons(*3*). In mammals, the macroglia is made up of radial glia-like cells (referred to as radial glia -RG-below), astrocytes, oligodendrocytes and ependymocytes. RG act as neural stem cells (NSC) and support adult neurogenesis, a process that promotes plasticity, growth and regenerative abilities in several species. The only glia in the nervous systems of early deuterostomes and early vertebrates were RG(*4*), but little is known about the actual heterogeneity of early glia and how glial diversity emerged. Additionally, the sequence of events of adult neurogenesis appears similar across vertebrates(*5*–*7*), yet a broad comparative analysis has never been conducted.

With its large RG population, abundant adult neurogenesis in regions homologous to mammalian neurogenic niches and intermediate glial diversity, the zebrafish telencephalon is an excellent model to address key questions of macroglia evolution and diversification(*8*). Here we used scRNA-seq to profile all cell types of the adult zebrafish telencephalon and generate an improved characterization of its macroglia. We then re-analyzed data covering the telencephalon from several vertebrates(*9*–*11*) which included glia (even if glia had been discarded in the original analyses), or the nervous system of invertebrates(*12*–*22*), and used these to determine to which extent the populations of RG and the neurogenic cascade were conserved and how glial diversity emerged from ancestral glia. These results improve our understanding of cell diversity in the zebrafish telencephalon, highlight conserved and divergent features of the neurogenic cascade across vertebrates, and reveal the multifunctional nature of ancestral astroglia.

### Generation of a molecular atlas of the adult zebrafish telencephalon

The zebrafish adult telencephalon is a major model for neuroscience studies, yet the limited characterization of its cell diversity hinders the labeling of specific cell types. Our cell collection procedure (Fig.1A) allowed us to enrich for RGs (located along the everted ventricular zone, green in Fig.1A) while still recovering the full extent of cell diversity in the adult zebrafish telencephalon. We first grouped cells into broad classes using well-defined markers (Fig.1B). We then subclustered each of those groups to identify refined cell subtypes. Because neurons and RG showed substantial heterogeneity we implemented a consensus-clustering strategy to resolve robust yet fine-grained clusters (Methods and Fig.S1A,B).

**Fig. 1.**
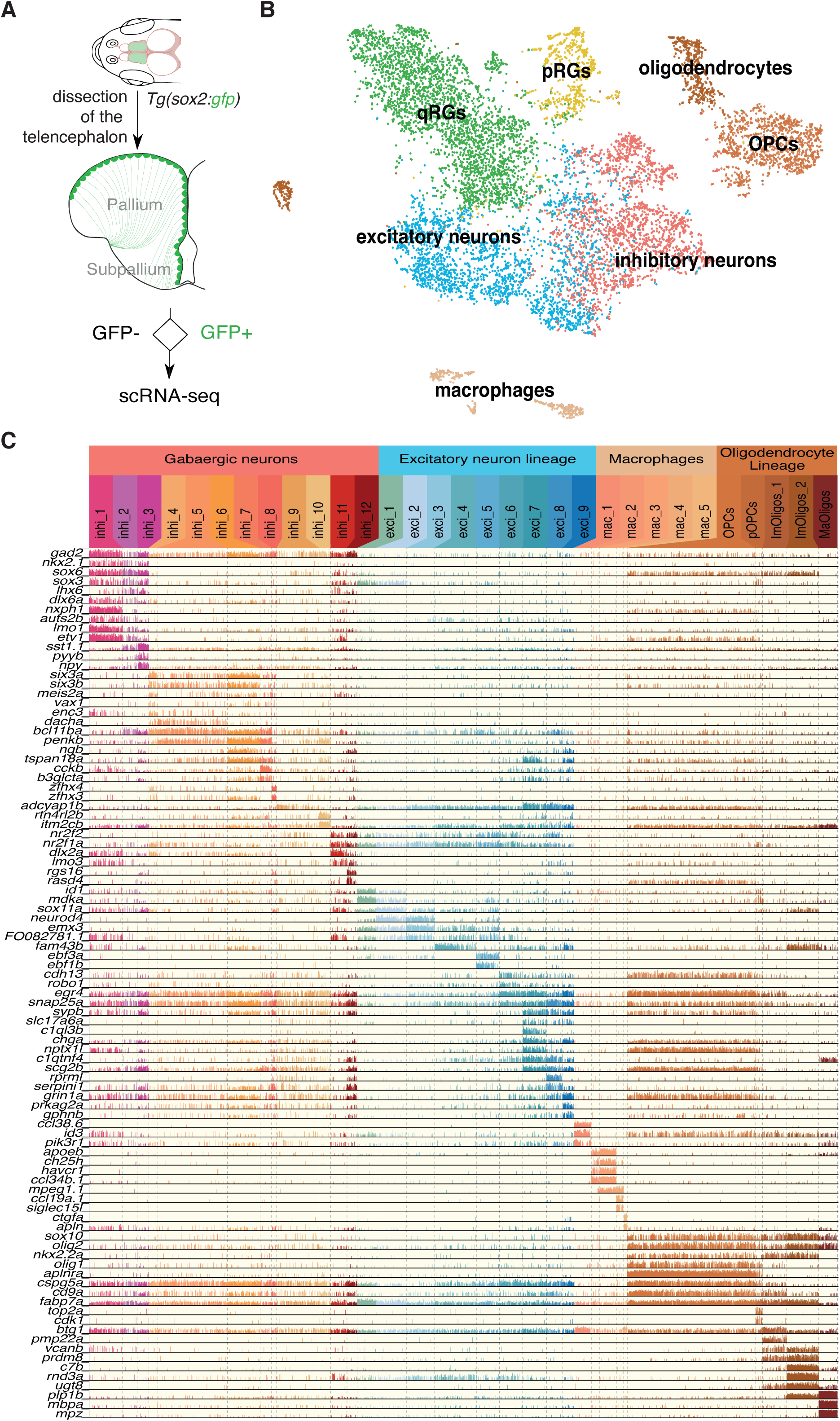
The adult zebrafish telencephalon is characterized by an extensive cell type diversity. **(A)** Cell collection strategy to enrich for RG. (**B)** tSNE of all cells colored by their broad cell type annotation. Abbreviations: q: quiescent, p: proliferating. (**C)** Expression of marker genes in refined non-RG cell clusters. Each color-coded column is a cluster and each line is a gene. Each bar represents one cell and the height of the bar reflects the level of expression of the gene.

This analysis first revealed notable results on non-RG cell types (Fig.1C). Among GABAergic neurons, we distinguished what likely corresponds to distinct ontogenies, compatible with an origin from homologs of the medial ganglionic eminence (MGE) (*nkx2*.*1*^*+*^, *sox6a*^*+*^), including somatostatin interneurons like in mammals, or the lateral ganglionic eminence (LGE) (*six3a*^*+*^, *six3b*^*+*^, *meis2a* ^*+*^) (Fig.1C). We also found that, based on their high expression of *nr2f1a* and *nr2f2*, a small proportion of GABAergic neurons likely derives from a homolog of the caudal ganglionic eminence (CGE), which has not been described in zebrafish before.

We also recovered unexpected diversity among brain immune cells despite a previous report suggesting that such heterogeneity was a human-specific trait(*23*). Among *mpeg1.1*+ cells we identified two clusters corresponding to functionally and ontogenetically distinct subpopulations of microglia previously described in the midbrain(*24*). We also identified another distinct and previously undescribed group of brain macrophages (Fig.1C).

### Ventricular patterning is conserved throughout life and evolution

We next turned to quiescent RG (qRG) (Fig.1B, green cluster). Conservative consensus-clustering followed by iterative cluster-merging based on differential expression identified 7 robust clusters (q1-q7) (Fig.2A, Fig.S1C). We further confirmed that this was not dictated by technical parameters and that clusters could be re-identified with a classifier (Fig.S2), and identified cluster-specific gene signatures (Fig.2B). There was little overlap between the clusters we identified and those proposed in a previous study(*25*). We thus reanalyzed the previously published data and performed in situ hybridizations (ISH) on whole mount telencephala and on serial coronal slices to assess the validity of our results. Although we resolved fewer clusters in these reanalyzed data(*25*) than in our own, likely due to the significantly lower number of cells profiled in this previous study, the data structure and patterns of cluster-specific gene expression now appeared consistent between the two datasets (Fig.S3). ISH further corroborated our clustering as genes enriched in the same cells showed similar patterns (Fig.S4).

**Fig. 2.**
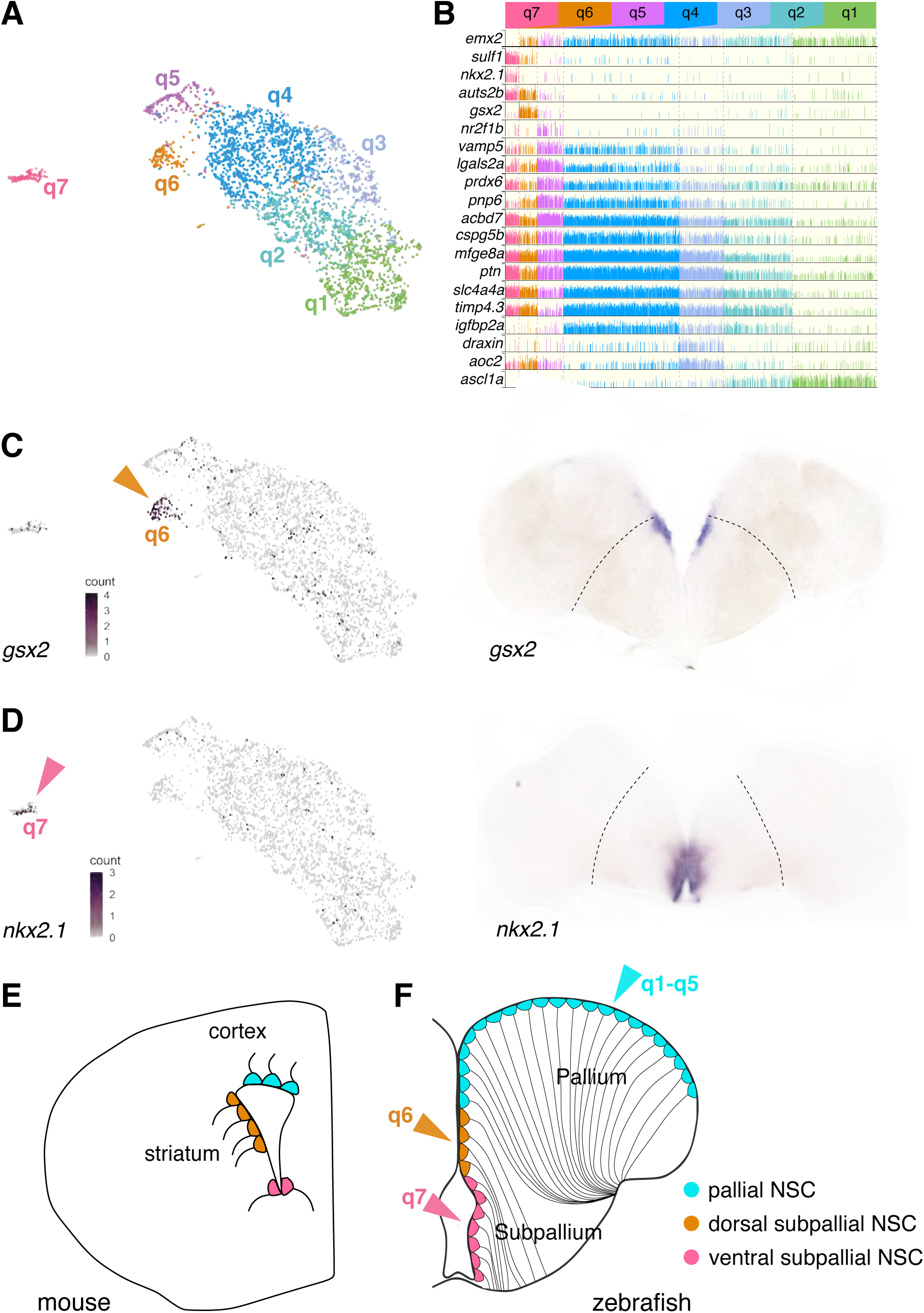
qRG in the zebrafish adult telencephalon are spatially patterned. (**A)** UMAP of qRG (green cluster from Fig.1B) colored by terminal cluster identity. (**B)** Expression of marker genes across qRG clusters, highlighting cluster-specific, regional and quiescence markers. (**C**,**D)** Orthologs of ganglionic eminences markers (*gsx2, nkx2.1*) projected on zebrafish UMAP (left, color-coded arrows matching (A) and (B)) and with ISH on coronal slices of the adult zebrafish telencephalon (right). Dashed lines indicate the boundary between pallium and subpallium. (**E)** Homologies of ventricular territories between zebrafish and mouse telencephalon (coronal sections) inferred from expression of regionalized transcription factor genes such as *emx2, gsx2* and *nkx2.1* (*10, 65, 66*). Depicted cells are RG, colored by their developmental origin (color-coded relative to clusters).

This analysis revealed spatially segregated RG populations in the ventricular zone. The *gsx2*+ q6 and the *nkx2.1*+ q7 clusters are restricted to a region close to the expected medial boundary between pallium and subpallium and in the ventral telencephalon respectively (Fig.2C,D). Similar observations were made in adult murine lateral ventricles, where the lateral wall expresses *Gsx2* and is LGE-derived, while the ventral wall expresses *Nkx2*.*1* and is MGE-derived(*10, 26, 27*). Markers for q5, including *nr2f1b*, were less strictly segregated but showed an enrichment caudally in pallial RG (Fig.S5) reminiscent of the *Nr2f1* gradient in RG of the developing mammalian neocortex(*28*).

Thus, ventricular progenitor patterning is maintained not only throughout life but also throughout evolution. In particular, the ventricular zone of the dorsomedial pallium (Dm) in zebrafish, which has been the focus of most of the studies on telencephalic neurogenesis in this model, is homologous to the dorsal wall of the SEZ in mouse. Conversely the *gsx2*+ area corresponds to the subpallial Vd domain rather than Dm, contrary to what was previously reported(*25*), and to the lateral wall of the mouse SEZ. These molecular landmarks (Fig.2E) will inform comparative studies of morphogenesis and neurogenic output between teleosts and mammals.

### Conserved and innovative features of adult neurogenesis

Adult neurogenesis has been mostly studied in rodents which display some of the highest evolutionary rates. Analyzing a broader range of species can improve generalizability and highlight fundamental features of the neurogenic cascade (*5*–*7*). By reanalyzing datasets published in amphibians(*11*), reptiles(*9, 29*) and mammals(*10, 30–35*) we compared gene expression patterns along successive steps of neurogenesis progression, from qRG to pre-activated RG (close to activation, corresponding to q1 in zebrafish) and neuroblasts, to reconstruct their state in the last common ancestors of different taxa (Fig.3). This revealed conserved expression of key regulators since the last common ancestor of osteichtyes. For example, the regulation of the Notch pathway is such that one Notch receptor gene is expressed in, and likely maintains, qRG(*36, 37*). Conversely, *Notch1* is expressed in paRG and so is the Notch ligand gene *DLL1*(*38, 39*). The *SoxC* genes *SOX4* and *SOX11* are turned on in paRG and reach their peak in neuroblasts, consistent with their described roles in hippocampal neurogenesis(*40*). Likewise, the EMT regulator *ZEB2*, which was recently found to be necessary for neurogenesis in the SEZ(*41*), displays conserved expression in neuroblasts. *SOX9*, which is a broad and specific marker of astroglia in adult mice(*42*) was not detected in previous datasets. We found that this is due to poor annotation of *sox9a* and that *SOX9* expression is conserved in qRG among osteichtyes.

Conversely, we found stepwise additions to the neurogenic cascade during mammalian evolution. ID transcription factors modulate the dynamics of Notch signaling. Among them *ID4* is more efficient than other IDs at inducing a return to quiescence(*43*), is the only one positively regulated by Notch signaling in qRG(*44*) and is a tetrapod addition (Fig.3). The switch from *NOTCH3* to *NOTCH2* in qRG likely happened early in the mammalian lineage, which, with the emergence of *NOTCH2NL genes* that potentiate NOTCH signaling in humans(*45, 46*), could explain some human features of adult neurogenesis (Supp. Text). We were unable to reconstruct neurogenic trajectories from large-brained animals(*33, 34*), possibly owing to sensitivity limits of untargeted scRNA-seq, although we confirmed evidence of ongoing adult neurogenesis when reanalyzing a large macaque dataset(*35*). While our re-analysis suggested that the initially defined RGL and IPC_2 clusters in this dataset are likely multiplets and of myeloid origin respectively (Fig.S6), these data did include progenitors not found in other datasets, allowing us to confirm the expression of some of the highly conserved genes in primates. In particular, *IGFBPL1* and *HES6* are highly and specifically expressed in IPCs and neuroblasts and thus represent promising markers to assess continued production of neurons in the human hippocampus (Fig.S6).

**Fig. 3.**
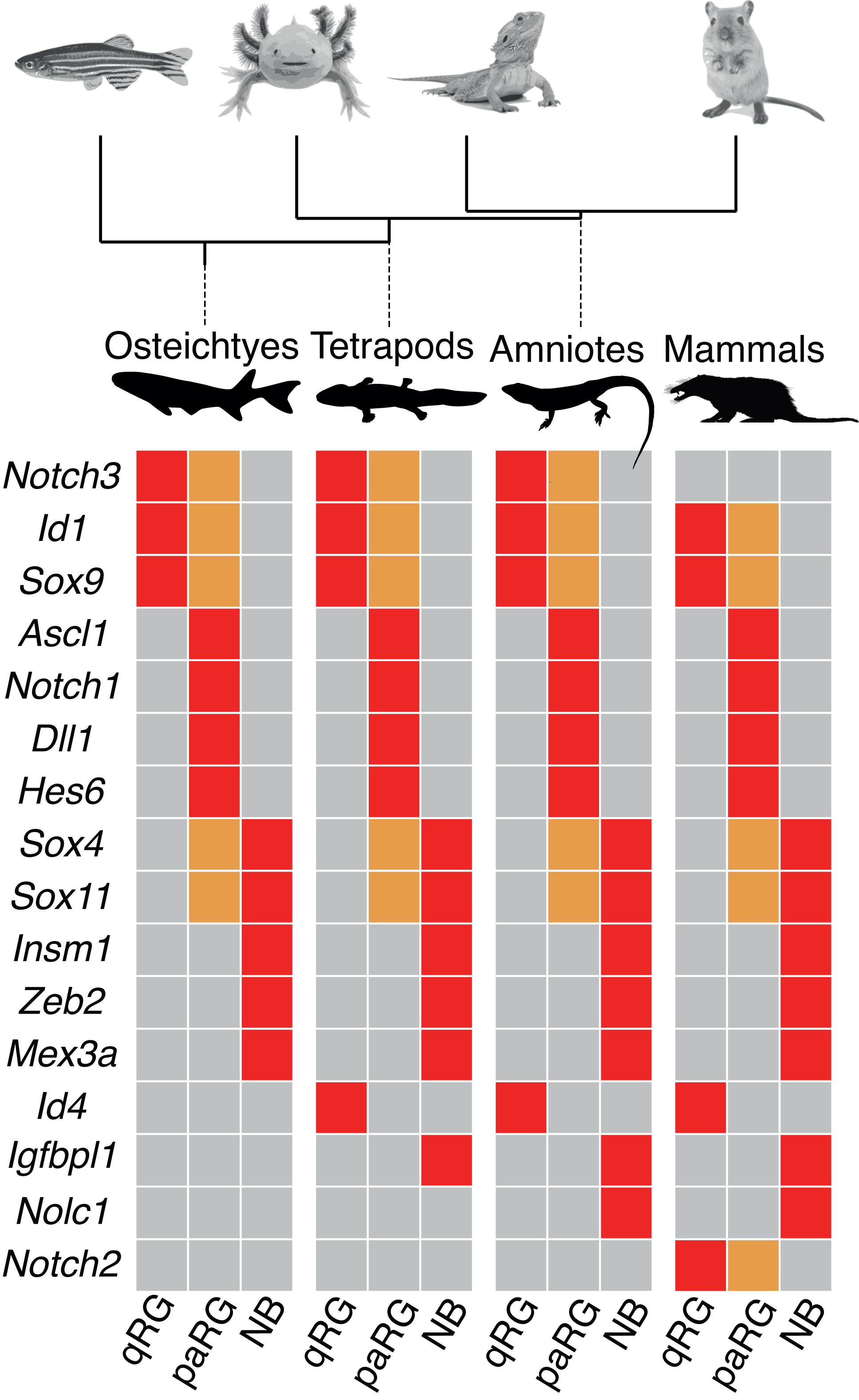
Conservation and variations in the evolution of the adult neurogenic cascade in vertebrates. Representation of the expression of putative key regulators of neurogenesis whose expression pattern differs across vertebrates. Top: phylogenetic tree depicting the profiled vertebrate species and the ancestors for which we reconstructed the expected pattern of expression when that applies. Bottom: trinarized relative expression depicted at different stages of progression along the neurogenic cascade (red: high expression, grey: no or low expression, orange: intermediate levels of expression). NB: newly born postmitotic neurons, paRG: pre-activated RG close to entering the cell cycle, qRG: quiescent PG

Together, this integrative approach highlights putative critical neurogenesis regulators either conserved throughout evolution or on the contrary responsible for taxon-specific properties.

### Evidence for quiescence depth heterogeneities in zebrafish qRG, and implications on the diversification of mammalian astroglia from ancestral RG

We found that our qRG clusters were further separated according to their quiescence depth (Fig.S7). Separation along a quiescence to activation trajectory was also apparent in mouse SEZ datasets. We focused on a recent one which profiled a large number of cells from the different regions of the SEZ with high sensitivity(*10*). To identify putative regulators of quiescence depth we developed a pseudo-ordering algorithm (Fig.S8A), applied it to both datasets and identified genes that show a strong association with deeper or shallower quiescence independently of region or species (Fig.S9). Some of these are likely part of a core quiescent stem cell gene set such as *HES1* which has been proposed to be a general marker of stemness(*47*), or those encoding the exosomal proteins CD9 and CD81 which are enriched in several types of quiescent cells(*48*).

Among the genes associated with deeper quiescence in zebrafish RG, several of them were expressed in both qRG and astrocytes in mice. We also noticed that some of these genes were instead highly enriched in, or even specific of astrocytes across several datasets (Fig.S10-S13) (*10, 30, 32, 49*). This came as a surprise since besides mammals, most species are not thought to have astrocytes(*50*–*52*). To further investigate the possible relationship between some qRG in zebrafish and mammalian astrocytes we focused on dorsal qRG (clusters q1 to q4, Fig.2A) to mitigate signals related to regionalization. We converted genes from zebrafish and mouse into a uniform nomenclature for orthologs (Methods) and mapped our pallial qRG to astroglia from mouse telencephalon (Fig.4A). This revealed that similarity with astrocytes in zebrafish RG increases with quiescence depth. Next, we scored mouse astroglial cells for genes enriched in q4 over q2 — two qRG clusters intermingled in Dm, with q4 predicted to be in a deeper state of quiescence— which revealed that astrocytes are enriched for q4 markers compared to RG (Fig.4B, Fig.S10-S13).

**Fig. 4.**
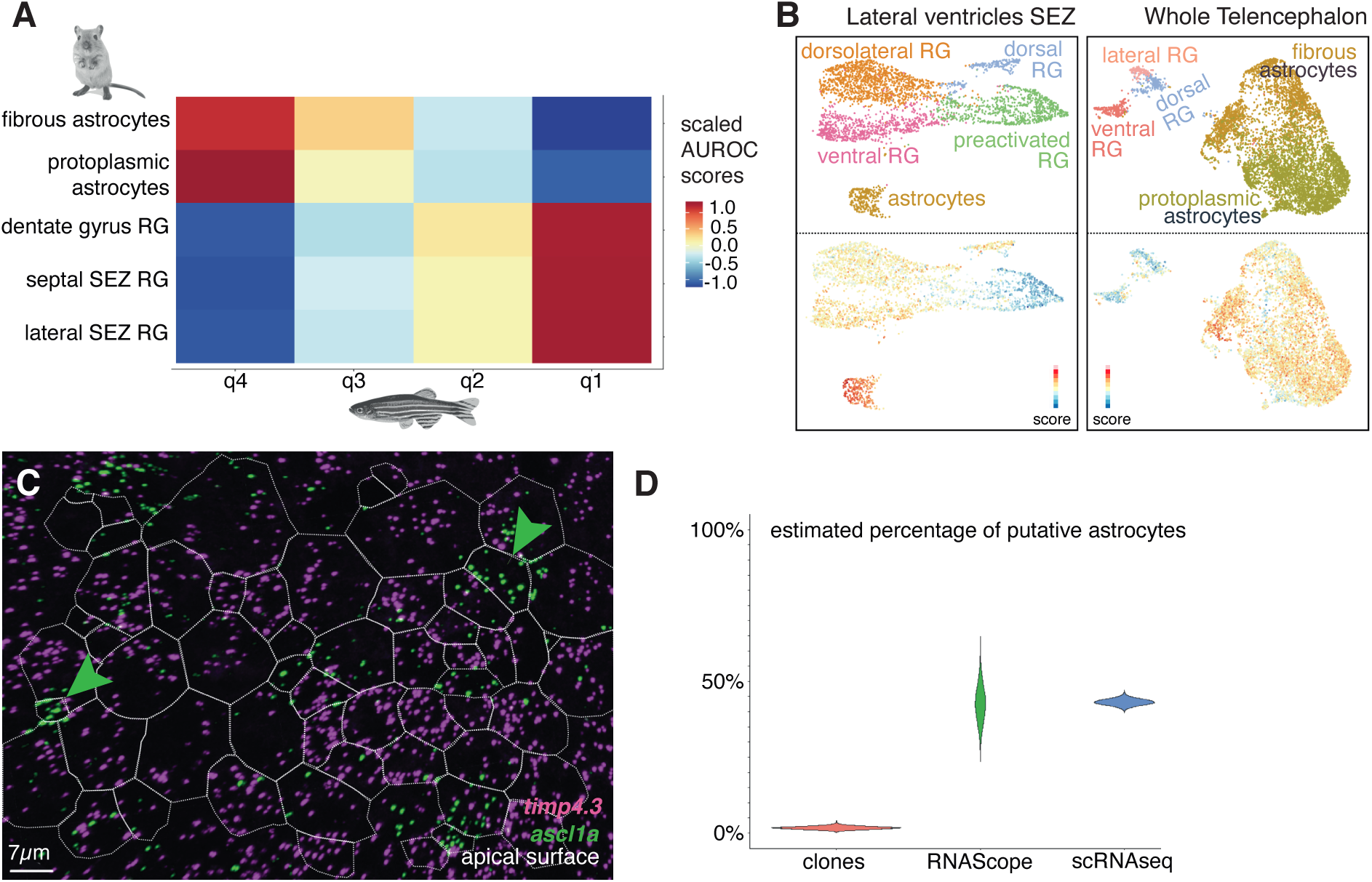
Homology between quiescent NSCs in zebrafish and mammalian astrocytes. **(A)** Cluster mapping between zebrafish dorsal RG (this study) and astroglia in the mouse telencephalon. (**B)** Scoring of mouse astroglial cells from (*10*) (left) and (*49*) (right) for genes enriched in zebrafish q4 over q2 revealing enrichment in astrocytes over RG. Top: annotated clustering of astroglial cells from each dataset. Bottom: Enrichment score for orthologs of genes enriched in zebrafish q4 over q2. (**C)** Representative image of double RNAScope ISH for *ascl1a* (green) and *timp4*.*3* (magenta) in zebrafish Dm (dorsal view), used to estimate the proportion of q4 cells (*timp4*.*3*^+^) in situ. ZO1 immunohistochemistry was used to delimit apical NSC surfaces (dotted lines). (**D)** Distribution estimations of the proportions of RG-only clones among those genetically induced in *Tg(her4:ERT2CreERT2)* adult fish in single RG then chased for 507 days (red), versus the proportions of q4 cells, measured either in situ with RNAScope (green) or in scRNAseq (blue). Estimations for the proportion of q4 cells are consistent and centered around 43%, while only 11 clones out of 630 exclusively contain RG cells after 507 days. The remaining 609 clones contain neurons, suggesting that most if not all Dm RG are neurogenic (p.value of two-sided binomial test <1.5e10-^132^).

Gene ontology analysis of the genes that were enriched both in q4 over q2 and in astrocytes over RG confirmed that this list of genes encodes proteins involved in astrocytic support functions such as neurotransmitter synthesis and recapture, metabolic support, maintenance of ionic balance and modulation of ECM properties (Table S1). Together with knowledge about the ontogeny and time of emergence of these cells, our results suggest that astrocytes evolved from ancestral RG through subfunctionalization in the mammalian lineage while astrocyte-like RG persist in zebrafish.

Next, we asked whether in zebrafish q4 astrocyte-like RG participate in adult neurogenesis or behave like radial astrocyte with no physiological NSC properties. The high level of transcriptome similarity among RG precluded lineage-tracing based on a specific q4 promoter. Instead, we analyzed the fate of clones from Dm genetically tagged with the Tg(*her4:ERT2CreERT2)* line (Fig.S14A)(*54*). *her4* is broadly expressed, enriched in q4 over q2 (Fig.S14B), and cells from q4 could be sorted from *Tg(her4:egfp)* reporter fish using the same *her4* promoter(*25*) (Fig.S14B). We reasoned that the ratio between the proportion of q4 cells in Dm and the proportion of *her4*-driven clones that do not produce neurons would allow us to infer whether q4 cells behave as NSC (Fig.S14C). At 507 days of chase, the proportion of RG-only clones was much lower than the proportion of q4 cells estimated in situ with RNAScope or in the scRNAseq dataset (Fig.4C-D, Fig.S14D,E), confirming that these cells do have a constitutive neurogenic potential.

Together these results suggest that mammalian astrocytes emerged from ancestral neurogenic RG that already expressed many of the genes related to astrocytic-specific functions and that persist as a deeply quiescent but physiologically neurogenic RG population in zebrafish. Importantly, our results also show that stem cells can perform differentiated cell functions when they are quiescent, contrary to the common view that quiescent stem cells are inactive.

### Evolution of astrocyte-like cells across planulozoa

Cell types similar to astrocytes appeared multiple times throughout evolution. Among vertebrates, the only class other than mammals where astrocytes are the major astroglial cell population are birds. Absence of astrocytes appears to be the basal state in reptiles, and turtles —the closest living relative to birds and crocodilians— do not have astrocytes either, suggesting that the cells described as astrocytes in birds evolved independently from those in mammals. We thus asked whether avian and mammalian astrocytes displayed a similar signature. We found that datasets generated from the high vocal center of zebra finch included both astrocytes and RG(*55*), with a substantial overlap between the top genes separating astrocytes from RG in birds and mammals (Fig.S15), suggesting that avian and mammalian astrocytes likely evolved in a similar way, from the same ancestral RG population.

The expression of a conserved astrocytic gene set in some zebrafish RG suggests that it emerged before the individualization of astrocytes as a cell type. Moreover, parenchymal glia associated with support functions are present in all major branches of bilaterians(*56*). To estimate the time of emergence of this astrocytic gene set we asked whether its genes were expressed across planulozoa by recovering and analyzing datasets from over 20 species(*9–22, 29, 34, 35, 49, 55, 57*), identifying existing orthologs and assessing their expression in glial clusters or among ectodermal cells (Fig.5). We found expression of the astrocytic gene set in all vertebrate RG, but not in Ciona ependymoglia (Fig.S16). On the other hand, we did not find any group of cells co-expressing genes from the astrocytic gene set in cnidarians or protostomes except in insects, where distinct glial subpopulations expressed part of the astrocytic gene set. Ensheathing glia, and to a lesser-extent astrocyte-like and perineurial glia, co-expressed several genes associated with mammalian astrocytes (Fig.S17). In addition, glial cells in protostomes did not express homologs of *SOX2* or *SOX9* genes, which are part of a conserved astroglial gene regulatory network in vertebrates.

**Fig. 5.**
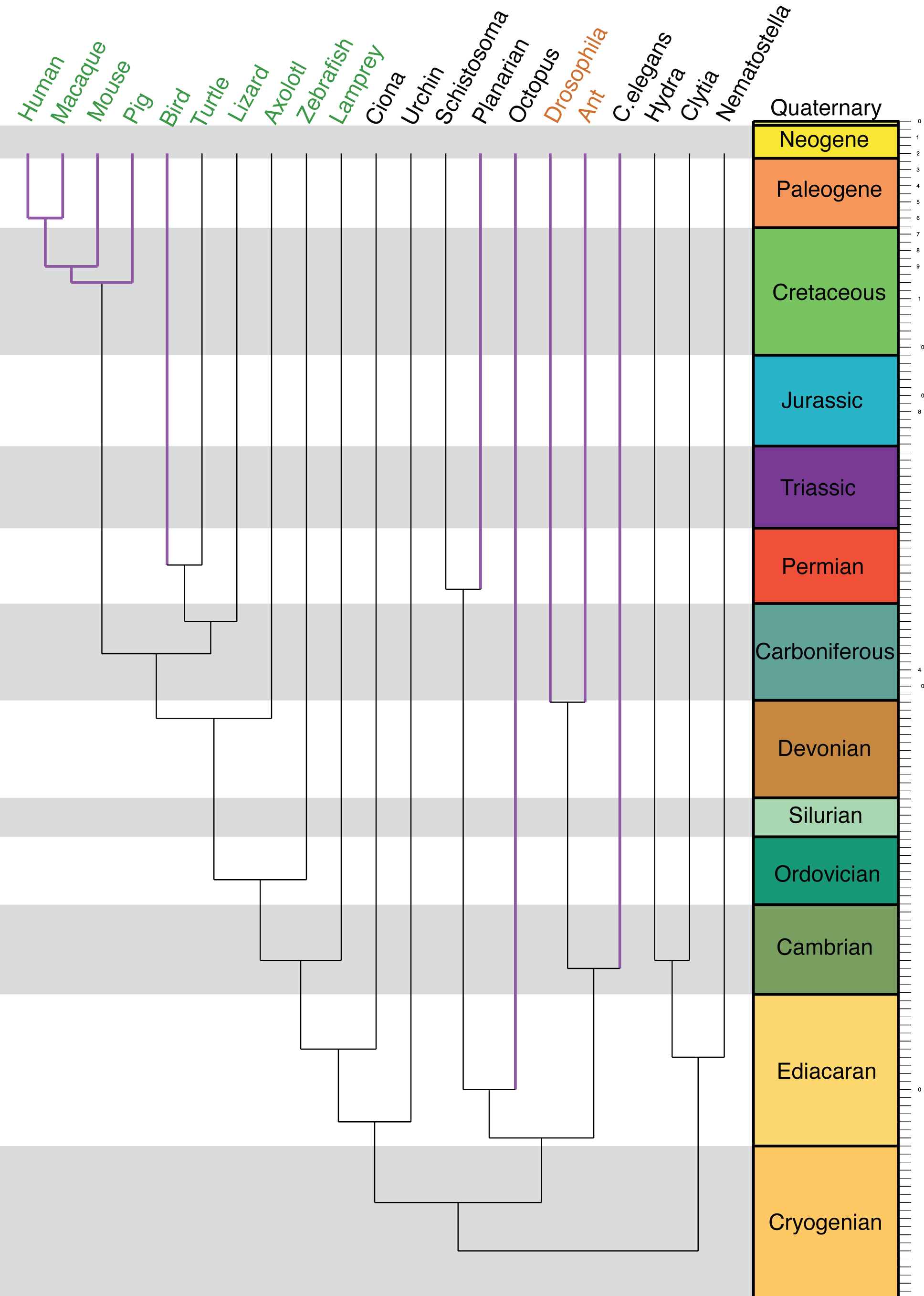
Emergence of the astrocytic synapomere. Phylogenetic tree depicting the expression of the astrocytic synapomere in analyzed species and whether parenchymal glia with supportive functions have been described in those species. Leaves in magenta represent phyla in which parenchymal glial cells have been previously described. Species in green co-express several genes of the astrocytic synapomere in the same glial cell clusters. Species in orange express several genes from the astrocytic synapomere but spread out across several glial cell clusters. Species in black do not seem to rely on the astrocytic synapomere.

This large-scale comparative analysis suggests that while avian astrocytes are sister cells to mammalian astrocytes, glial cells described in protostomes are not. Several genes involved in astrocytic functions started being expressed in vertebrate RG and make up an astrocytic synapomere(*58*). The emergence of parenchymal glia in insects, which together express many of the genes of the astrocytic synapomere, is likely the result of homoplasy, suggests remarkable functional convergence and highlights the interest of studying aspects of glial physiology in protostomes.

### Evolution of other macroglial cells

Cells of the oligodendrocytic lineage and ependymocytes make up the macroglia alongside astroglia. We used our zebrafish telencephalon dataset to identify cells belonging to the oligodendrocytic lineage and compared expression of putative core regulators and effector genes across chordates (Fig.S18). Others recently proposed that cells expressing *sox9b* are zebrafish astrocytes(*59*). We found that they are transcriptionally closer to cells of the oligodendrocytic lineage and express several genes associated with oligodendrocyte maturation such as *c7b, rnd3a, ugt8, cadm4, gpr17, myrf* and *plp1b*, as well as low levels of *mbpb*. These cells might thus correspond to maturing oligodendrocytes, similarly to a population in the larval spinal cord(*60*). For inter-species comparisons, we grouped cells based on the expression of *CSPG4* (and/or *PDGFRA* for tetrapods). A core set of transcription factors made up of *NKX2*.*2, SOX10, OLIG2* and *OLIG1* is largely conserved across vertebrates. *NKX2*.*2* and *SOX10* are also expressed in lamprey glia, although orthologs for *olig* genes have not been identified in the lamprey genome. *CSPG4* (or *NG2*) which is commonly used to label OPCs is also widely conserved across vertebrates and detected in lamprey glia, contrary to *PDGFRA* and *APLNRA* which are restricted to tetrapod and zebrafish OPC respectively. The expression of genes involved in myelin formation appears variable, with the exception of *MBP* and *PLP1*. In scRNA-seq studies, only low levels of *plp1a* were detected whereas a recent proteomic study found that PLP1B is 86 times more abundant(*61*). We found that this discrepancy results from an incomplete gene model for *plp1b* which is indeed expressed at high levels in zebrafish oligodendrocytes. Lamprey glia express not only *PLP1* but also *MPZ*, like jawed fish, suggesting that co-expression is the ancestral state, lost in tetrapods. Despite being commonly associated exclusively with PNS, myelin *PMP22* was detected in most species besides mouse. *PMP22* being restricted to the PNS is thus a derived trait in mice. The core gene set conserved across vertebrates is not present in Ciona suggesting that the emergence of glia with oligodendrocytic properties happened in vertebrates after they split from other chordates.

Ependymocytes likely appeared as a cell-type later in vertebrate evolution. In the zebrafish telencephalon similar cells are restricted to a rostral and ventral area(*62*) and were not profiled in our dataset. We detected a few in lizard and birds(*9, 55*) where they have been observed in small numbers along the ventricles whereas in mammals they cover the whole ventricular surface and represent a much larger fraction of astroependymoglial cells in the niche. Ependymocytes express several markers of qRG, but simultaneously display remarkable convergence with other multiciliated cells which appear to have co-opted a similar multiciliated cell apomere (Fig.S19A). We found that mouse ependymocytes express regulators of SEZ neurogenesis that are not detected in lizards or birds, suggesting that they acquired additional regulatory roles in mammals (Fig.S19B).

## Discussion

Our comprehensive molecular and cellular characterization of the adult zebrafish telencephalon expands knowledge on telencephalic evolution and will facilitate targeted studies of its diverse cell populations. In particular, we identify underappreciated heterogeneity among brain macrophages and conserved ontogenies of interneuron families across vertebrates.

In addition, using a novel consensus-clustering approach we identified distinct robust subpopulations of qRG. This is independent of the specific genome assembly, feature choice or clustering parameters and is reproducible across datasets and supported by subsequent cross-species comparisons and in situ validation. Heterogeneity among RG is explained to some extent by spatial origin, with homologous territories between mouse and zebrafish maintaining their positional identity from development to adulthood.

The presence of a caudo-rostral gradient of *nr2f1b* shows that this gradient predates neocortex arealization. The organization of the zebrafish pallium is less well-defined than that of the neocortex and appears to have evolved independently(*63*), suggesting that *NR2F1* might have been co-opted to specify sensory areas(*28*) after the split between actinopterygians and sarcopterygians. Of note, expression of *wnt3a* has been shown to be restricted to the lateral pallium in zebrafish(*64*). Therefore, both caudo-rostral and medio-lateral gradients involved in hippocampus specification were already present in the last common bony fish ancestor.

Comparisons of adult neurogenesis across vertebrates revealed a highly conserved neurogenic cascade. Besides generating a list of putative critical regulators of adult neurogenesis in vertebrates, our approach highlights core principles of neurogenesis shared across phyla. Stepwise additions to this conserved core occurred during evolution and likely underlie specific properties of adult neurogenesis and NSC maintenance in different species, which can now be tested via anachronistic gene expression manipulation. Likewise, a switch from *NOTCH3* to *NOTCH2* in mammalian qRG, together with the emergence of functional *NOTCH2NL* genes in humans, provides a realistic and testable hypothesis for the premature depletion of RG in humans (Supp. Text). We also identify genes that show a conserved association with quiescence depth, including some showing consistent patterns in other tissues and/or cancers. Further investigation of those candidates is thus likely to yield fundamental insights on the maintenance of stemness and quiescence.

Our data suggest that avian and mammalian astrocytes evolved from the same ancestral multifunctional RG and that similar RG persist in zebrafish. The synapomere associated with those sister cell types is enriched in genes involved in metabolism, neurotransmission fine-tuning and extracellular milieu homeostasis. Several of these genes can also be detected in lamprey glia but not in putative Ciona ependymoglia. Likewise, the genes that segregate in oligodendrocytes in gnathostomes and are expressed in lamprey glia can induce intricate membrane re-organization and prevent axonal degeneration. Together, these observations are consistent with an ancestral role in enabling neuron communication and survival via the establishment of structures that later served as templates for the development of other functions such as myelination with which we usually associate macroglia. Macroglia then diversified through subfunctionalization, division of labor, co-option of new modules and neofunctionalization.

We expected to detect cells that retained expression of a similar gene set across bilaterians but were unable to do so in non-deuterostomes besides insects. Although we attempted to maximize sensitivity by surveying many datasets and including only those with high coverage, this approach can produce false negatives for rare cell populations or due to imperfect genomic annotations. Alternatively, it is possible that the functions associated with the astrocytic synapomere were initially performed by other cells including neurons themselves and became the property of glial cells in vertebrates. In that case, even if a common set of ancestral glia existed before they might not have expressed genes enriched in current vertebrate macroglia and would have been missed with our approach. Additional targeted studies will be helpful to shed light on the origins and evolution of glia, their coevolution with neurons and will benefit from focusing on genes identified in this study.

Finally, the presence of astrocytic-like RG in vertebrates that do not have *bona fide* astrocytes, such as the zebrafish, has several implications, including on the notions of quiescence and dedifferentiation (Supp. Text). Moreover, it raises interesting questions regarding the evolution and other potential functions of these cells. Are they also neurogenic in salamanders and reptiles? How do they behave upon injury and to what extent do they contribute to regeneration? Understanding the behavior of these cells, which look like intermediates between RG and astrocytes, could facilitate reinstatement of stem cell properties in astrocytes to promote regeneration.

## Acknowledgments

We thank the ZEN team for input, the Institut Pasteur CB UTechS service platform for expert assistance with FACS analysis and scRNAseq processing, and Thibaut Brunet for critical reading of the manuscript. We are greatly indebted to Isabel Fariñas for sharing data prior to publication, and for insightful discussions.

## Author contributions

Conceptualization: DM, LBC; Methodology: DM, IF, AA; Funding acquisition: LBC; Project administration: LBC; Supervision: LBC; Writing – original draft: DM; Writing – review & editing: DM, LBC.

## Data availability

All data used in the analysis are available at: https://www.ncbi.nlm.nih.gov/geo/query/acc.cgi?acc=GSE225863

## Supplementary Materials

Materials and Methods

Supplementary Text

Figs. S1 to S19

Table S1

Supplementary references (*67*–*165*, in gray in Main text’s list)

