## Supplemental material for "Integrative single-cell transcriptomics clarifies adult neurogenesis and macroglia evolution"

**The PDF file includes:**

Materials and Methods
Supplementary Text
Figs. S1 to S19
Table S1
References (*67–165*, in gray in Main text)

### Materials and Methods

Fish lines and maintenance

All procedures relating to zebrafish (*Danio rerio*) care and treatment conformed to the directive 2010/63/EU of the European Parliament and of the council of the European Union. Zebrafish were kept in 3.5-liter tanks at a maximal density of five per litter, in 28.5°C and pH 7.4 water. They were maintained on a 14-hours light/10-hours dark cycle (light was on from 8 a.m. to 10 p.m.) and fed three times a day with rotifers until 14 days post-fertilization and with standard commercial dry food (GEMMA Micro from Skretting) afterwards. All fish used for experiments were between 3 and 4 months old. Fish from the transgenic line *Tg(sox2::GFP)* were maintained on an AB background.

Euthanasia

Fish were euthanized in ice-cold water (temperature comprised between 1° and 2°C) for 10 min, according to a special dispensation and following the guidelines of the Ministry of Superior Education, Research, and Innovation.

Dissociation and cell sorting

Cell sorting was conducted three days in a row to collect replicates, using twenty 3 months old adults from the *Tg(sox2::GFP) line* (*67*) on each day. Brains were dissected in Ringer's solution. The telencephalon was separated from the midbrain and the olfactory bulbs were removed. The two hemispheres were separated and cut along the boundary between pallium and subpallium to enrich for pallial cells. Cell dissociation was carried out according to (*68*). Cells were then sorted on a FACSAria III. We used forward and side scatter and DAPI staining to distinguish live cells from debris and sorted cells on their GFP levels to enrich for *sox2*-expressing cells while still including GFP negative cells to not miss any relevant population. After sorting, encapsulation of cells and reverse transcription was immediately performed using the 10x Chromium Controller and Chromium Single Cell 3' Kit v2 and then immediately frozen at -80°C until all replicates had been collected.

Library preparation and sequencing

After reverse transcription all replicates were processed in parallel with the v2 Kit. After barcoding libraries were pooled and split over multiple lines of a HiSeqX and sequenced at a depth over 100k reads per cell using 2x150 paired-end kits with the following sequencing read recommendation Number of Cycles: 26 cycles Read1 for cell barcode and UMI, 8 cycles I7 index for sample index and 98 cycles Read 2 for the transcript. This yielded a saturation above 95% for all libraries.

Mapping and filtering of data

Initial analysis was conducted using the cellranger software. Reads were demultiplexed and mapped to the GRCz11 zebrafish genome assembly from Ensembl, although we later replicated our analyses with a genome assembly curated by the Lawson lab (*69*). Datasets were first analyzed individually to determine whether they were fit for integration. First we filtered cells based on the number of genes (nGene) and UMIs (nUMI) detected as well as on the relationship between nGene and nUMI. nGene is expected to be positively correlated with nUMI. Lower nGene than expected for a given nUMI value suggests low library complexity whereas higher nGene than expected for a given nUMI value suggests excessive library complexity and likely doublets. To filter on that criterion, we fit a loess regression curve for nGene~nUMI with a span of 0.5 and of gaussian family and removed cells which residuals were beyond three standards deviation of the mean (*70*). Then we inspected the frequency of cells associated with a given nGene. This yielded a distribution with a narrow peak at low levels of nGene followed by a broader distribution centered around a peak close to nGene=900. We considered the first peak to be low quality cells and set a threshold to remove it from the rest, which ended up being nGene=200. We also removed cells which had abnormally high nGene or nUMI based on the distribution on a plot of nGene as a function of nUMI. Then we removed genes expressed in fewer than 10 cells. Finally, we computed the percentage of the transcriptome that consisted of mitochondrial gene (percent.mito) and inspected plots of percent.mito as a function of nGene. These two variables show negative correlation because high percent.mito tend to be observed in low quality cells, in which fewer genes are detected. We set a threshold at 10% which removed a tail of cells with high percent.mito and low nGene. Subsequent inspection of the variation of gene expression as a function of technical variables such as nGene after regular scaling and normalization revealed that variance in gene expression was not dependent on technical factors. This pre-integration analysis also suggested that the second replicate was of lower quality than the other two which was consistent with the fact that it took longer to go from cell sorting to cell capture. Thus we only used replicates 1 and 3 for subclustering analyses to avoid clusters driven by technical artifacts.

Identification of clusters and their markers

We did not need to use any batch removal method to obtain good merging between the replicates. Variable genes were identified by selecting genes exhibiting high variability given their level of expression without using a fixed general threshold. We selected PCA components to include based on whether they explained over 1% of the variance across the first 100 principal components and whether that variance was flagged as significant by the JackStraw test (*71*). Initial clustering was performed on all the cells using a simple Smart Local Moving algorithm (*72*). We looked for and removed doublet clusters using three separate approaches. The doublet cells methodology from the scran package (*73*) was used to score individual cells and the doublet cluster methodology to detect clusters which look like a mixture of two other clusters. An adapted DoubletFinder algorithm was used to generate doublets from randomly selected pairs of cells and detect cells which frequently co-clustered with those mock doublets (*74*). This multi-pronged approach ensured that we did not keep any artefactual cluster at this stage.

Several broad cell types were already subdivided into multiple clusters. We isolated each individual lineage and performed further clustering to identify fine-grained cell subpopulations. Substantial heterogeneity was apparent in neurons and RG. To ensure that we did not overcluster we developed a consensus clustering approach. We pooled results from Bayesian model mixture, smart local moving algorithm, multilevel algorithm, walktrap, spinglass and density peaks. kNN graphs used for the graph clustering approaches were themselves built using edges weighed either as in SNN-Cliq (*75*) or Phenograph (*76*). From this we obtained a consensus matrix with the Cluster-based Similarity Partitioning Algorithm (*77*). From this consensus matrix we drew a final hierarchical clustering and used a conservative cutoff to identify robust clusters. We then subjected each pair of neighbouring clusters to differential gene expression detection to determine whether they should be merged. On the final partition, we used hypergate to detect combinations of genes that best identify each cluster (*78*). We then used these lists to train a classifier and assess its performance, taking an area under the curve above 0.7 as indication that clusters could be re-identified with a limited number of genes.

The approach with hypergate as well as regular differential gene expression analysis using MAST with nUMI as a variable to regress against yielded lists of markers for each clusters.

Gene Set Enrichment Analysis

GSEA requires ordered lists of genes rather than restricted lists of markers (*79*). We used each gene to train a classifier to recover given clusters and used the area under the receiver-operator curve as the value for gene set enrichment analysis. The area under the curve is a bounded value between 1 and -1 and its sign indicates whether increased expression of the gene improves recovery of the cluster (positive sign) or removal of the cluster (negative signe), thus its properties are particularly suitable to this type of analysis. We recovered gene sets from Zfin and carried out the analysis using the liger package https://cran.r-project.org/web/packages/liger/ (not to be mistaken with rliger which is dedicated to dataset integration (*80*)).

Trajectory reconstruction

To identify step-wise transitions in expression from deep to shallow quiescence we performed a pseudo-ordering analysis with a custom algorithm inspired by a method developed by the Satija lab (*70*). We first embedded cells in a neighbourhood-preserving reduced space using diffusion maps on variable genes as implemented via the diffusionmap R package (https://cran.r-project.org/web/packages/diffusionMap/index.html). In this reduced space we computed a thousand minimum spanning trees using two thirds of the data and calculated the average distances between cells along bootstrapped minimum spanning trees. From this distance matrix we used multidimensional scaling to compute a three-dimensional embedding allowing good visualization of the reconstruction. In their implementation, Mayer et al then finalized the reconstruction with a final minimum spanning tree (*70*). However, previous studies showed that parametrizing a minimum spanning tree with a skeleton of nodes improves reconstruction and avoids short-circuits (*81*). We thus used the ElPiGraph R package to fit a skeleton of up to a hundred nodes (*82*). Noisy paths can form due to uneven density in the embedding, we thus pruned independent paths that emerged and terminated in the same cluster. We then completed the tree and projected each cell on its edges. Branches were then automatic

Synthesis of digoxigenin probes for in situ hybridizations

Total cDNA was synthesized from brain lysates through reverse transcription with Superscript II. Genes of interest were amplified via polymerase chain reaction with specific primer pairs derived from https://primer3.ut.ee/ and/or https://www.ncbi.nlm.nih.gov/tools/primer-blast/ and cloned with the help of the StrataClone PCR Cloning kit from Agilent. Genes that could not be cloned this way were ordered as gblock from IDT https://eu.idtdna.com/pages/products/genes-and-gene-fragments/double-stranded-dna-fragments/gblocks-gene-fragments with overhangs allowing restriction digest cloning into NEB10 bacteria (#C3019H). 5µg of DNA was linearized with the appropriate enzyme, purified and transcribed in vitro in presence of digoxigenin-labeled uridine and precipitated with lithium chloride. Probes were then resuspended in 50µL DNAse and RNAse-free water and stored at -80°C.

Preparation of tissue for immunohistochemistry and in situ hybridization

Whole brains or whole telencephala were dissected from 3 to 4 months old fish after euthanasia as described above in cold PBS. They were immediately placed in cold 4% PFA and fixed on a rotating platform at 4°C overnight. The next day brains were dehydrated through sequential washes in 25%, 50%, 75% and finally 100% methanol (mixed with PBS the first three solutions) and then stored at -20°C. For labelling, brains were first rehydrated through washes in 75%, 50% and 25% methanol in PBS and then processed with the appropriate method.

Chromogenic in situ hybridizations

Adult brains were retrieved from -20°C and rehydrated. They were subsequently treated with a bleaching solution made up of 0.5X SSC, 3% H202, 0.05% formamide diluted in DNAse and RNAse-free water. They were then washed 4 times in PBS + 0.1% Tween 20 and then pre-incubated at 65°C in a solution containing 5X SSC, 65% formamide, 0.1% Tween 20, 50µg/µL of heparin and 2.5mM of citric acid for at least 4 hours. The solution was removed and then added again with the labeled antisense probe for overnight incubation at 65°C. The next day the probe was removed and the brains washed in increasingly stringent conditions up to 0.05X SSC to remove excess probe before being incubated in blocking buffer with an anti-digoxigenin antibody coupled to horseradish peroxidase (HRP) for 2 hours. The brains were then washed overnight and the next day HRP activity was revealed with NBT/BCIP either on whole mounts or on 60µm thick coronal slices prepared after the antibody step.

Single molecule FISH with RNAScope

We sought to leverage the increased sensitivity of single molecule fish and its ease of coupling with immunostaining against proteins to better quantify the presence of specific cells and in particular to better distinguish between cells expressing high or low levels of *timp4.3.* The initial steps up to bleaching are the same as those performed for regular chromogenic ISH. Then, we pre-incubated brains at 40°C either in RNAScope's proprietary diluent buffer from Bio-Techne (https://www.bio-techne.com/reagents/rnascope-ish-technology) or in chromogenic ISH buffer but with 25% formamide instead of 65%. Hybridization was performed in the same conditions as pre-incubation and we followed instruction from the Hiplex kit for the rest of the procedure.

Clonal analysis

Data for clonal analysis was reanalyzed from (*54*). Traced fish were double transgenic generated from crossing Tg*(her4.1:*ERT2CreERT2)/+ (*83*) and *Tg(-3.5ubb*:loxP-EGFP-loxP-mCherry*)*/+ (*84*). Irreversible expression of mCherry was sparsely induced by injections of low tamoxifen quantities at 3 months and fish were traced for up to 507 days. Cell types were identified on the basis of the expression of glial and proliferation markers as well as their position relative to the ventricle. For more information see (*54*). Because glia and in particular astrocytes can proliferate in mammals, we considered all clones which included only RG, whether one or many, as potentially originating from non-neurogenic cells and compared their proportion with that of cluster 4 cells.

Re-analysis of published datasets

We downloaded raw matrices for around 40 datasets for re-analysis (see text for references). We subjected them to the same type of quality filtering and analysis as our own datasets without the consensus clustering approach. In most cases we were interested in synexpression of specific gene-sets independent of precise clustering. In the case of the Cebrian-Silla dataset (*10*) we used the gene sets identified in the paper to assign regional identity to the different clusters of neural stem cells. For the whole mouse telencephalon dataset (*49*) our own analysis agreed almost perfectly with the published annotations but we found that regional signatures from (*10*) were differentially enriched in subsets of the original "SVZRG" cluster and thus subdivided it further. Astrocytes were also heterogeneous but we could not assign a specific identity to astrocytic subsets and thus kept the broad annotations.

Identification of orthologous genes lists

To compare lists of enriched genes between species and to perform direct mapping of data sets we needed reliable lists of orthologs. We found that commonly used methods limit the feasibility of this type of comparison. A widespread approach is to convert all of the genes in a dataset to their orthologs through Ensembl (https://www.ensembl.org/index.html ) and to then take all one to one orthologs for integrated analysis. We found that annotation through Ensembl makes many mistakes, both false positive and false negative. Moreover because teleosts underwent an additional round of whole genome duplication compared to other vertebrates, many mammalian genes map to multiple orthologs in zebrafish which leads to them being discarded from such analyses. We thus curated a lists of orthologs based on a common syntax originating from the one used by eggnog (*85*). We identified orthologs for zebrafish genes by integrating information from eggNOG, Alliance Genome (https://www.alliancegenome.org/), Ensembl and Zfin (https://zfin.org/) both through command line tools and manual curation of identified relationships. We mapped mouse genes to the same set of orthologs in a similar manner but without input from Zfin. The common syntax being based around eggNOG's output then allowed us to directly compare genes with orthologs mapped through eggNOG for other datasets for example in birds or lizards. We also mapped genes from less studied species through eggNOG and Ensembl Metazoa to assess the expression of individual genes in those datasets.

Cross-species mapping

To map our zebrafish data to the one generated by (*49*) we used our curated lists of orthologs for both mouse and zebrafish, generated matrices with the same gene names and aggregated them based on genes that were expressed in both datasets. We identified variable genes using the variance stabilizing transformation (*86*) and used those as input to MetaNeighbour (*53*) to unbiasedly identify similar cell types between the two species.

### Supplementary Text

Homology of neurogenic niches

Regionalization of the SEZ niche in the mouse brain has been the object of study for many years (*65*, *90–98*). The mouse SEZ can be subdivided into several microdomains with distinct molecular signatures, correlated to the type of progeny the stem cells can produce and to the developmental domain they originate from. In particular, distinct domains of the pallium and subpallium give rise to specific regions along the dorsoventral axis of the adult neurogenic niche, with neural stem cells retaining expression of transcription factors from the developmental period to adulthood. Adult neurogenesis in the zebrafish telencephalon has been almost exclusively studied in the dorsomedial palium (Dm region) (*99*). Although anatomical subdivisions of the telencephalon have been proposed based on the layout of neurons (*99*), no direct relationships linking the different ventricular territories between the everted telencephalon of actinopterygians and the evaginated telencephalon of most other vertebrates had been established.

Our work shows that major dorsoventral subdivisions along the telencephalic ventricle in zebrafish mirror those found in mammals, and in particular that Dm is likely homologous to the dorsal wall of the lateral ventricle which harbors deeply quiescent neural stem cells in the mouse (*96*, *98*). This corrects a previously proposed territorial subdivision of the telencephalic ventricles (*25*) and lays the groundwork for further comparisons across vertebrates. For example, RG in Dm proliferate less than RG from the *gsx2* domain in the adult zebrafish telencephalon (*100*). This mirrors the relative levels of proliferation between the dorsal and lateral walls of the ventricles in adult mouse and suggests that functional correlates of positional identity are also conserved to some extent. In addition, although it has been shown that a structure reminiscent of the rostral migratory stream exists in the zebrafish telencephalon (*101*), whether the neurons that are generated are similar to the newborn olfactory neurons in mammals and whether different regions of the ventricle contribute in the same way is unknown. Further studies, combining molecular phenotyping and lineage tracing in zebrafish while leveraging knowledge from the results obtained in mammals, could shed light on these questions and highlight potential adaptations related to adult neurogenesis that happened over the course of evolution to improve olfaction.

We did not find support in favor of the existence of a population homologous to dentate gyrus granule cells in our data. This could be due in part to a limitation of our approach which aimed at enriching for glia and thus likely underrepresents the full diversity of neuronal populations in the zebrafish telencephalon. Although the extra-ventricular position of the hippocampus appears to be mammalian-specific, recent data confirms that the different hippocampal subfields are also individualized in reptiles (*29*) . However, in urodele amphibians this appears to not be the case (*11*)and instead neurons co-expressing CA and dentate gyrus markers are found in the medial pallium (*102*). Nonetheless, our results and previous studies (*103*) show that gradients of expression involved in specifying the hippocampal pallium are present and functional investigations have identified a region of the caudal dorsolateral zebrafish pallium that is involved in spatial encoding (*104*, *105*). Thus it is uncertain how closely stem cells from the hippocampal pallium might be related between species and to what extent this might be correlated with cell type diversification in the mammalian medial pallium. We could not identify a subpopulation of cells from the *wnt3a*+ domain previously observed with in situ hybridizations (*103*), likely because these cells are very few in number and might also have been selected against during our dissection. A medio-lateral gradient of expression of morphogens in RG was described in axolotl with molecular signatures reminiscent of those in mammals and reliably delineating different regions of the pallium(*11*). Similarly, in our re-analysis of *Pogona vitticeps* data (*106*), we found that a subset of RG expressed high levels of *WNT3A* which displays medial pallium expression in mammals. Unfortunately, neither *WNT3A* nor *PROX1* were reliably detected in a pioneering spatial transcriptomic approach on *Pogona vitticeps* telencephalon (*107*), which prevented us from confirming that the *WNT3A*-expressing RG are in the same territory as the dentate gyrus-like neurons.

We anticipate that these remaining questions will be addressed soon through larger datasets on teleost brains and targeted studies across vertebrates which will shed light on the origin of cell type diversification in the hippocampus. This is of particular interest for the study of adult neurogenesis as the dentate gyrus niche has been described as being more of a mammalian innovation (*108*) whereas the SEZ would be more of a vestigial trait, despite the fact that the architecture of the mammalian SEZ is also noticeably different from the ventricular zones of non-mammalian vertebrates. A greater understanding in the molecular and lineage structure similarities across vertebrate will be necessary to substantiate or definitively invalidate such a hypothesis.

<u>A mechanistic hypothesis on the rarefaction of adult neural stem cells in humans</u>

The existence of adult neurogenesis in humans has been a highly debated topic since it was first hypothesized. In the last few years several studies investigating human neurogenesis have reported contradicting results and reignited the debate (*109–116*). Single-cell RNAseq datasets from the human and non-human primate hippocampus have been generated with some claiming to detect neural stem cells. We reanalyzed these datasets (*33–35*, *117*, *118*) but we were unable to reconstruct a complete neurogenic cascade in humans or macaque. In particular our reanalysis is consistent with another report (*114*) that concluded that cells initially described as neural stem cells (*117*) were in fact likely ependymocytes. Thus, although there is some evidence that immature neurons can still be detected in adult humans, adult neural stem cells remain elusive and it is more largely agreed that they are likely prematurely depleted compared to most other species (*119*). However, we lack a conceptual framework to explain how and why such a situation might have arisen despite the advantages of ongoing adult neurogenesis with respect to functional plasticity and regenerative abilities. Our analysis of the molecular cascade involved in neurogenesis regulation highlighted the important role of the Notch pathway. Strikingly, the main Notch receptor expressed in quiescent RG is different in mammals compared to lizards, amphibians and fish, suggesting that a switch between paralog resulted in *NOTCH2* taking up the functions of *NOTCH3* in mammals. Such switches between paralogs are common during evolution(*120*). It was recently shown that human-specific genes derived from duplication of the

*NOTCH2* locus –the *NOTCH2NL* genes– potentiate regular Notch signaling and promote symmetric divisions in RG early on, which ultimately results in an enlarged cortex (*121*, *122*). Other studies also showed that potentiating Notch signaling could result in longer maintenance of RG early on, but that this is later followed by an increased terminal differentiation into astrocytes (*123*). Taking all this into account, the switch between *Notch3* and *Notch2* in mammals, leading to durable co-expression of *NOTCH2* and *NOTCH2NL* genes in human RG (*121*, *124*), could lead to accelerated RG depletion and the lack of RG-like cells in adult humans. This model would explain why this change is specific of humans while other non-human primates, which do not have functional *NOTCH2NL* genes, retain adult neurogenesis, as has been demonstrated by radiographic, histologic and sequencing studies (*125–127*, *35*). Contrary to arguments explaining the low neurogenesis in humans based on a selection against plasticity due to the size and complexity of their brain(*128*), this interpretation suggests a tradeoff with an expanded neocortex. Selection for these traits would have taken place at a moment when sight was already the primary sensory modality thus reducing selective pressure on olfactory processing (*129*), and when life expectancy was much shorter, thus resulting in only a short period of time without hippocampal neurogenesis given that it persists well into teenage years. Most importantly, this model can be experimentally tested. First, several experiments modulating the Notch pathway have been conducted with results supporting the proposed hypothesis (*44*, *123*, *130–134*). Second, copy number variations in the *NOTCH2NL* locus are a well-known cause of neurodevelopmental disorders leading to the creation of patient cohorts (*121*, *124*). Using current work aimed at developing ways of quantifying the presence of stem cells in vivo (*113*, *135*), or post-mortem histological studies on tissue from patients with fewer *NOTCH2NL* copies, will make it possible to eventually found further correlates in support of our model directly in humans.

<u>Emergence and diversification of glia</u>

Relatively little is known about the evolutionary origins of glia (*136*). Several phyla appear to contain representatives both with, or apparently devoid of, glial cells. Moreover, glial cells are much more diverse than neurons in appearance, molecular profile and functions. Whether glia emerged or not alongside neurons is disputed and what they emerged from is unknown. It has been proposed that, because neurons are very demanding energetically and inefficient at controlling the extracellular changes caused by their activity, glial cells taking up these functions must have emerged at the same time as neurons in epithelial nerve nets (*137*). However, not only do some species appear to not have any glial cells, but in some species with glial cells, like *C. elegans*, these are not necessary for survival (*138*, *139*). Moreover, neurons can perform functions often associated with glial cells, for example in copepod where conduction speed is increased by a sheath derived from neurons themselves (*140*). Our comparative analysis also revealed cases (e.g., in Ciona, Fig.S16) where genes usually associated with glial functions appear expressed at high levels in neurons instead. We do not believe that our approach is powerful enough to confirm that neurons in species with a limited glia complement are less dependent on support cells. However, in light of descriptions from the literature and our observations, we do believe that functionally investigating this hypothesis is worthwhile, for example by assessing Cnidarian neurons' abilities to recapture neurotransmitters and provide their own source of energy. It is apparent that neurons and glia have not evolved separately but rather have co-evolved, and it would be interesting to determine whether this has resulted in neurons becoming less self-sufficient in presence of glia.

The astroglial system of actinopterygii has been described as having undergone only moderate evolutionary modifications, and they are generally thought to lack bona fide astrocytes (*50*). This implied that astrocytic functions were either unnecessary or fulfilled by another cell type, with RG as the most likely candidate to do so (*141*). Although recent work showed that hindbrain RG in zebrafish were capable of acquiring intricate morphologies (*142*) and communicating with neurons (*143*), these abilities are shared between astrocytes and RG-like cells in mammals (*144–151*). Thus, while these results highlight fundamental properties of RG, they cannot by themselves confirm the hypothesis that zebrafish RG can behave like astrocytes.

Here, we found that distinctions among RG were partially driven by a group of genes which also separates astrocytes from RG-like cells in mammals. Within the limits of scRNAseq and quantifications using a few markers with RNAScope, we almost never observe proliferating or *ascl1a*+ cells that express high levels of astrocytic markers, although clonal analysis suggests that all RG in the zebrafish pallium eventually give rise to neurons. This suggests that the majority of astrocytic-like cells recovered in the scRNAseq data correspond to a substate of quiescence for RG. This implies that their ability to fully perform support functions is coupled to state transitions: astrocytic genes are expressed at high levels when the cells are deeply quiescent and turned off as they activate. This suggests that before subfunctionalization and the emergence of distinct cell types fulfilling those roles in terrestrial vertebrates, ancestral astroglial cells underwent a form of temporal cell differentiation allowing them to perform both functions but not simultaneously. This is reminiscent of the current model of metazoan evolution based on observations of unicellular organisms capable of colony formation (*152*, *153*). This sequential dimension differs from previous examples of cell-type evolution described in the brain with ancestral neurons that simultaneously co-expressed sets of genes that became segregated after subfunctionalization (*29*). This also has important implications for our understanding of quiescence and differentiation. It shows that contrary to a popular representation the quiescence state is not —or at least not necessarily— a state of rest but can rather be an opportunity for a stem cell to fulfill functions not related to the production of progeny. Moreover, if cells physiologically oscillate between a state where they are ready to activate and produce progeny and a state in which they are quiescent and performing functions usually associated with differentiated cells, we must be very careful in defining cases of regenerative de-differentiation.

Our approach does not allow us to pinpoint the time when cells emerged that display permanent astrocytic functions and no physiological ability to generate neurons. Such cells could have first appeared without losing their contacts with the ventricles and be present in some of the data we have analyzed. However, without a reliable quantification of the proportion of RG capable of generating neurons, we cannot support or invalidate this possibility. Delamination of astroglia to give rise to parenchymal astrocytes appears to be linked to parenchymal thickness. It has been hypothesized that brain enlargement leads to stretching of the radial process which itself causes delamination (*154*). Work from the Kalmán lab, in particular on crocodilians, suggests that the number of parenchymal astrocytes in species with low numbers of them is not directly correlated to thickness (*155*). However, close crocodilian relatives with thicker brains show that the enlarged areas are areas that likely contained comparatively higher numbers of astrocytes in archosaurian ancestors. This suggests that parenchymal astrocytes can appear without a substantial thickening of the parenchyma and then engage in a self-reinforcing loop. On one hand the presence of dedicated parenchymal astrocytes which can more efficiently fulfill support functions might be beneficial to cope with increased metabolic needs (*156*) and relax constraints on parenchymal thickness. On the other hand the increase in thickness itself can promote more astroglial delamination to give rise to parenchymal astrocytes.

Independently of whether parenchymal astrocytes precede or follow parenchymal enlargment, a large parenchyma is associated with large numbers of parenchymal astrocytes and with reduced

RG numbers. Birds are remarkable in this regard, as they have very thick brain parenchyma and a large number of astrocytes, yet also retain large numbers of RG acting as neural stem cells comparatively to mammals, squalomorphs (*157*) or hagfish (*158*). It would thus be interesting to turn back to birds, which played a major role in the early days of the study of adult neurogenesis (*159*, *160*), to better understand how the balance between astrocytic differentiation and self-renewal of neural stem cells can be regulated. Further comparative studies across species harboring parenchymal astroglia will also allow us to determine the level of similarity between these cells and their generative process. This would in turn shed light on whether the differentiation landscape of RG is limited. Hagfish and platypus represent two highly interesting unconventional models to characterize in this context. Recent work on lamprey suggests that their RG co-express astroglial and oligodendroglial genes (*57*) and that they exhibit morphological traits consistent with a dual function (*161*). Assuming that this is the ancestral vertebrate state, the nature and behavior of hagfish parenchymal glia are highly intriguing and they might represent yet another sister cell type to other glia with features of both astrocytes and oligodendrocytes. Similarly, one study reported that contrary to other mammals, monotremes appear to not have distinct oligodendrocytes and astroglial cells but rather a hybrid cell type (*162*) (although these results have never been corroborated to our knowledge). Given efforts to improve the usefulness of hagfish (*163*, *164*) as a model and with single-cell transcriptomics having already been applied to platypus (*165*) it is likely that in the near future studies will answer these questions and further improve the resolution of the phylogenetic tree of glial cell types.

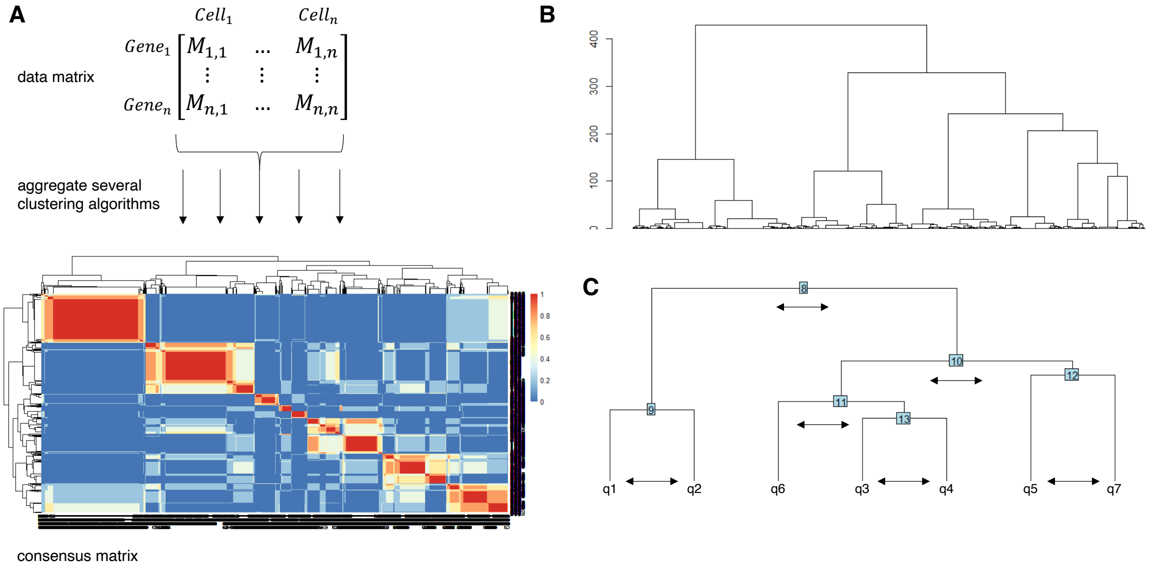


**Fig.S1. Approach used for clustering quiescent radial glia. (A)** Schematic of the consensus clustering approach. (**B)** Dendrogram obtained from hierarchical clustering on the final consensus matrix for the entire dataset. (**C)** Dendrogram of final qNSC clusters, arrows depict the combinations that would be subjected to differential gene expression analysis to determine whether they need to be merged or not.

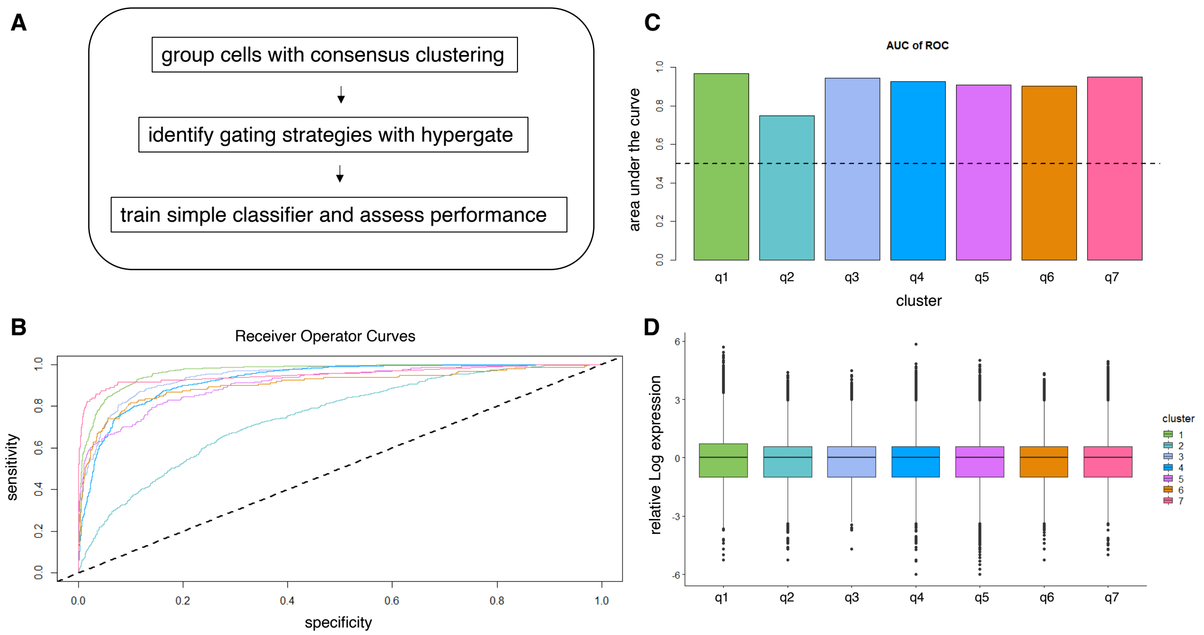


**Fig.S2. Verification of the validity of clustering**. (**A**) Schematic of the approach used to verify cluster actionability, inspired by Valentine Svensson (https://www.nxn.se/valent/2018/3/5/actionable-scrna-seq-clusters). (**B**) Receiver Operator Curves depicting how well clusters are recovered with the unfiltered output from hypergate, dashed line represents performance expected by chance, line color matches cluster colors used in other figures. (**C**) Area under the Receiver Operator Curves from B, dashed line represents performance expected by chance. All clusters get an AUC over 0.7. q2 likely being in an intermediate stage between q4, q3 and q1, was expected to get the lowest performance. (**D**) Relative Log Expression for all genes across all clusters. The distributions are centered around 0 in all cases suggesting that clustering is not driven by cell quality.

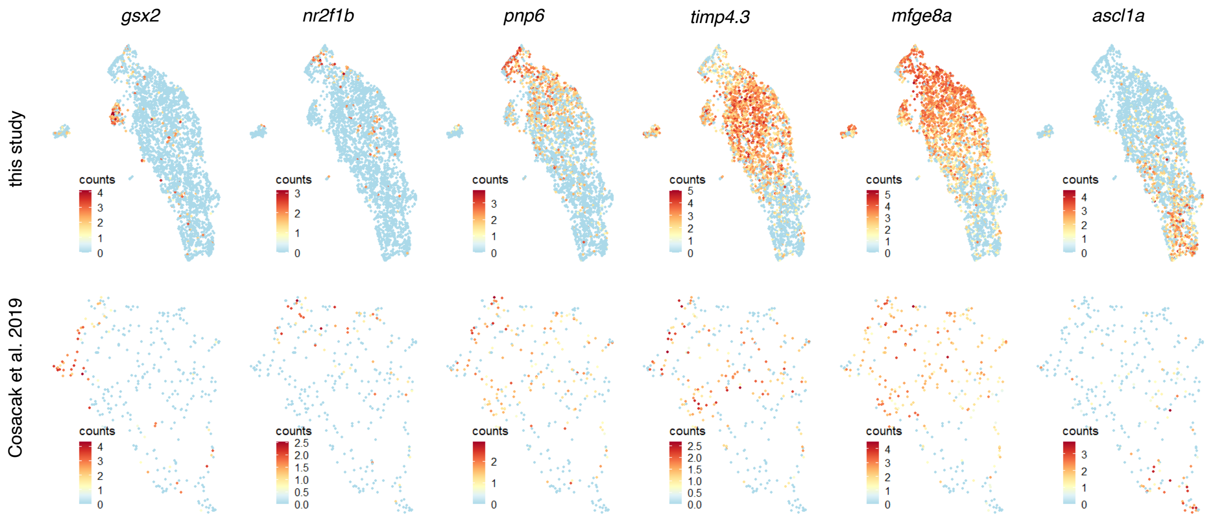


**Fig.S3. UMAP expression of cluster markers in qRG**. Expression of genes enriched in different clusters in our dataset and comparison with a previously published dataset that profiled 10 times fewer cells (*25*), here reanalyzed with our clustering method (Fig.S1). Gene expression patterns are quite similar in both datasets and consistent with our proposed grouping of qRGs rather than the one proposed in (*25*).

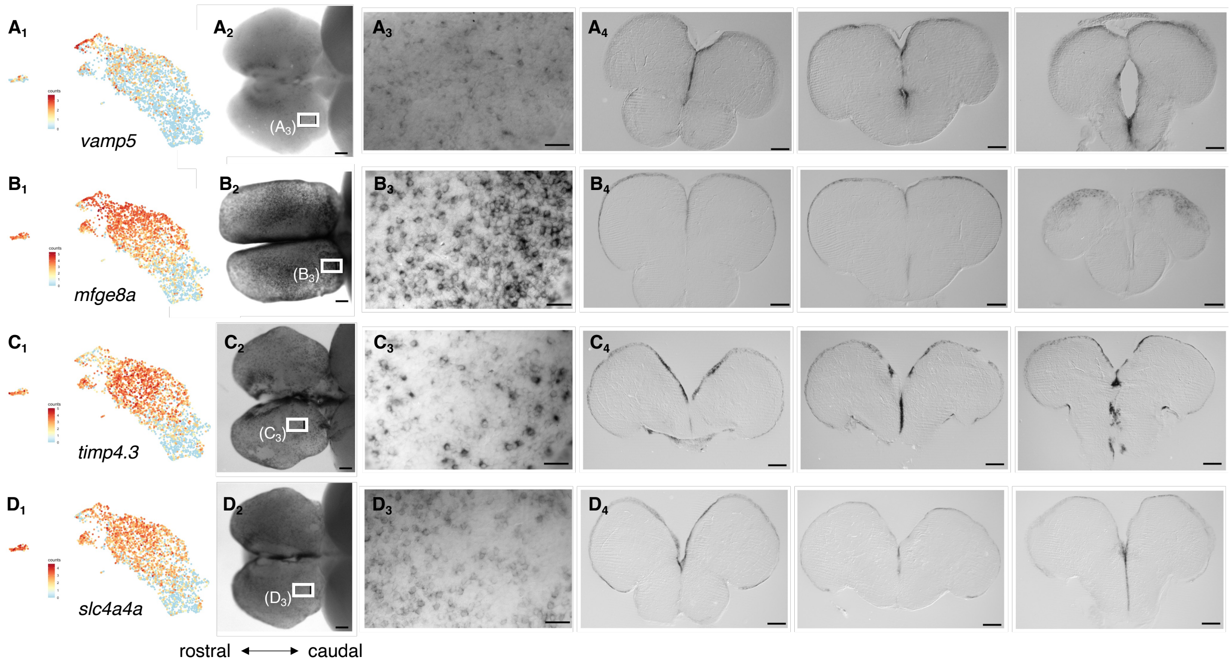


**Fig.S4. In situ validation of genes that show variable expression among qRG in the zebrafish adult pallium**. Chromogenic ISH validates the relative extent of expression of genes enriched in subsets of qRGs and confirms the existence of distinct qRG populations intermingled with each other in the zebrafish adult pallium. Each line follows the same pattern with gene expression plotted on the qRG UMAP, a whole mount image in dorsal view, a close-up of a region in the caudal part of the dorsal pallium (boxed on the whole mount image) and three coronal slices at different positions along the rostro-caudal axis from rostral to caudal. Scale bars≈500µm.

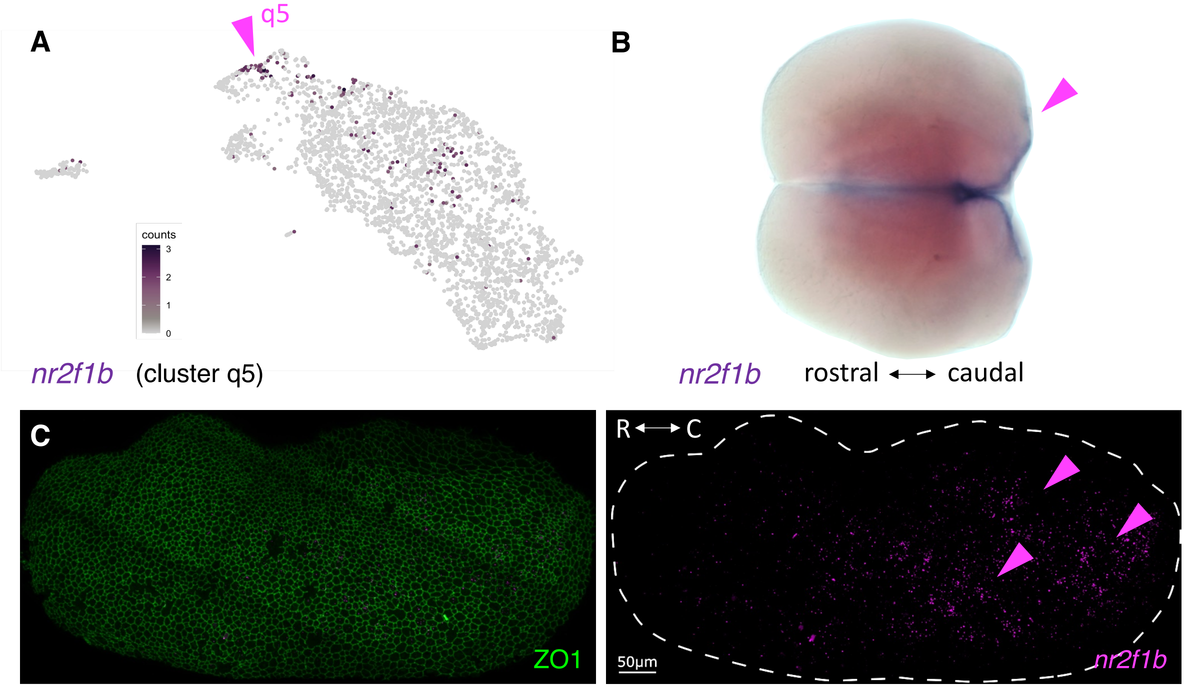


**Fig.S5. Caudo-rostral gradient of *nr2f1b* expression in the zebrafish adult pallium**. (**A)** *nr2f1b* expression projected on the qRG UMAP, showing specificity of expression in q5 (arrowhead, see also Fig.2B). (**B)** Dorsal view of a zebrafish telencephalon stained for *nr2f1b* expression by chromogenic ISH. (**C)** Dorsal view of a hemisphere of a zebrafish telencephalon (confocal microscopy, z=20µm projection), double stained in whole-mount for ZO1 (green, immunohistochemistry) and *nr2f1b* expression (magenta, RNAScope ISH). The two channels are shown separately to appreciate the caudo-rostral gradient of *nr2f1b* expression.

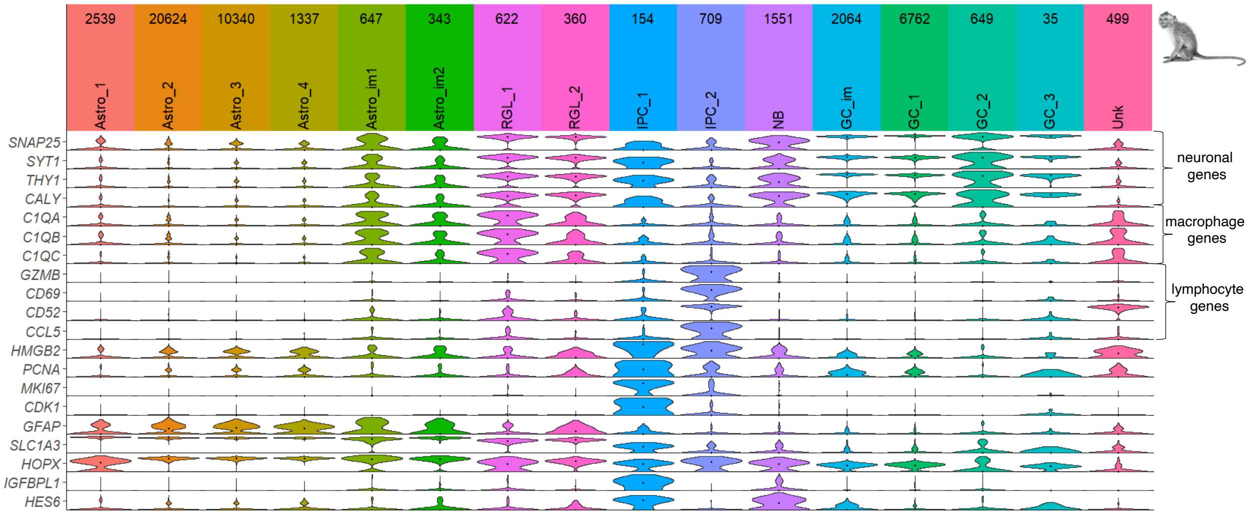


**Fig.S6. Re-analysis of Macaque dentate gyrus scRNAseq** (*35*)**.** Violin Plot for selected genes in an adult macaque hippocampus dataset with the initially proposed cluster identities (re-clustering gave comparable results). We included astrocytic clusters as well as clusters proposed to be involved in adult neurogenesis. *HOPX* is one of the genes most enriched in RG-like cells (RGL) over astrocytes in mice but is expressed at lower levels in putative macaque RGL clusters which also express several genes supposedly specific of neurons or macrophages suggesting that those are multiplets. Putative IPC_2 expresses several lymphocyte-specific genes and is likely a population of cycling blood cells. IPC_1 expresses intermediate levels of astrocytic and neuronal genes and high expression of cell-cycle related genes, consistent with its annotation. *IGFBPL1* and *HES6* show high expression in IPC_1 and negligible expression in clusters other than IPC_1 and neuroblasts, making them valuable candidates to distinguish between continued neurogenesis or persistence of DCX in developmentally-born neurons in humans.

Astro: Astrocytes, GC: granule cells, IPC: intermediate progenitor cells, NB: neuroblasts, RGL: radial glia-like cells, Unk: unknown. Im: immature

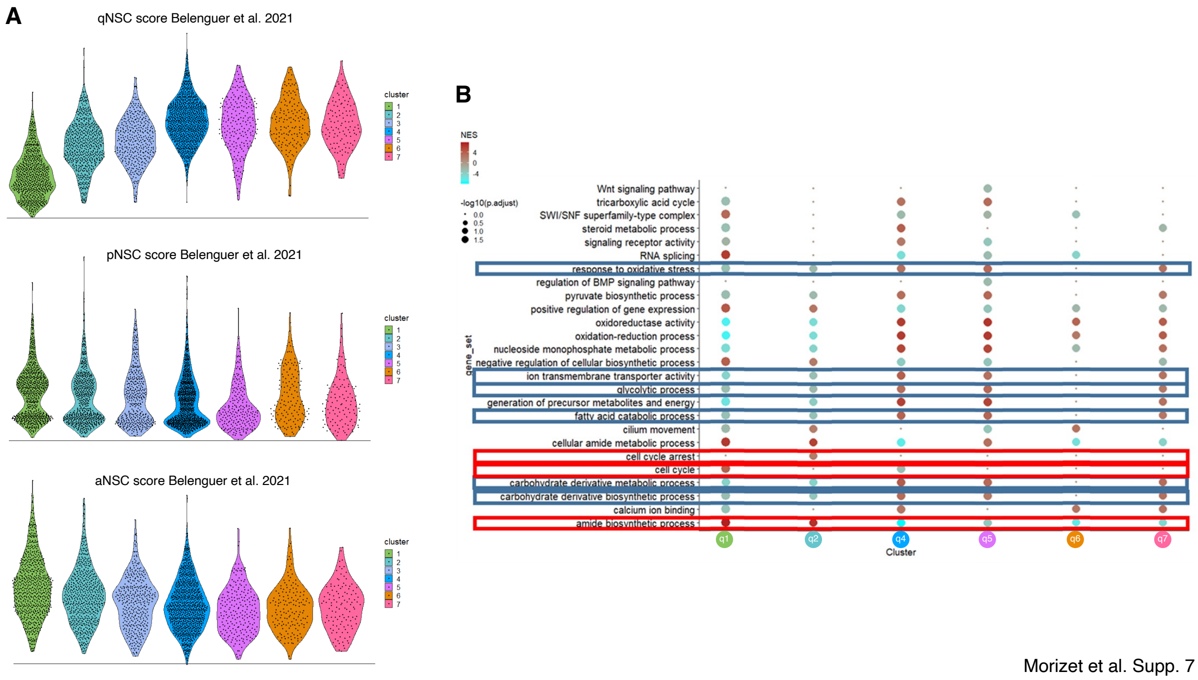


Morizet et al. Supp. 7

**Fig.S7. qRG are in different depths of quiescence**. (**A**) Violin plot depicting, in zebrafish telencephalic qRG clusters, the expression levels of genes the orthologs of which are transcriptionally associated with quiescent, primed and activated RG (respectively labeled qNSC, pSNC and aNSC) in mice (dataset derived from (*87*)). Note the graded representation of q1 to q4 signatures along the a→p→q progression. (**B**) Gene Set Enrichment Analysis on zebrafish qRG clusters. Gene sets highlighted in blue and red are associated with quiescence and activation, respectively. One column for each cluster, except for q3, which was too close to q4 to identify specific gene sets and is not depicted individually here.

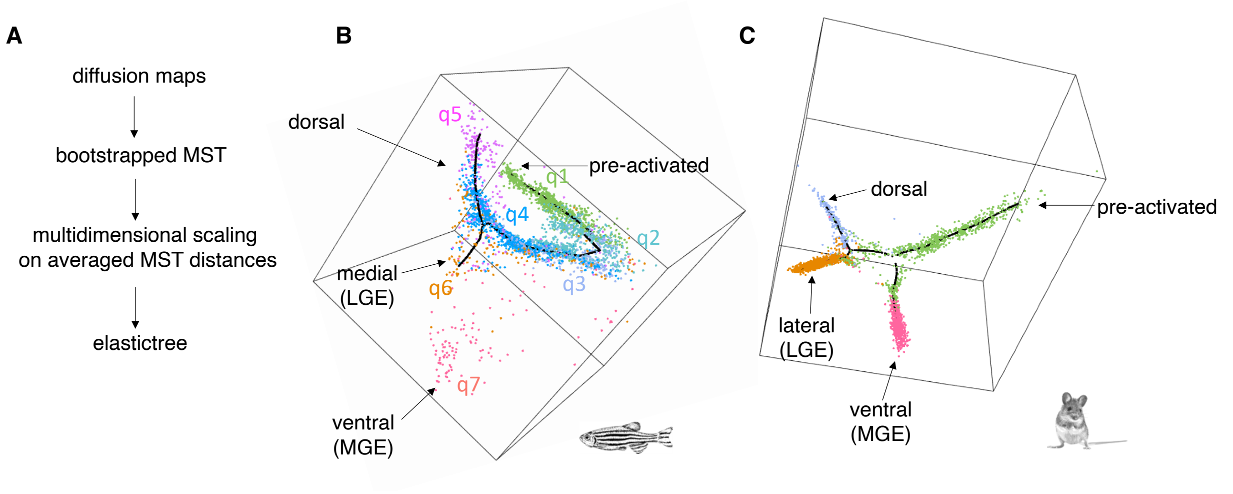


**Fig.S8. Pseudo-ordering reconstruction in zebrafish and mouse.** (**A**) Pipeline developed to determine pseudo-ordering, inspired by (*70*) and (*81*). (**B**) Annotated pseudo-ordering of qRG in the zebrafish adult telencephalon, color-coded as in Fig.2A. (**C**) Annotated pseudo-ordering of qRG from the mouse SEZ from (*10*), colors are matched to those of zebrafish for the same regions and for pre-activated cells.

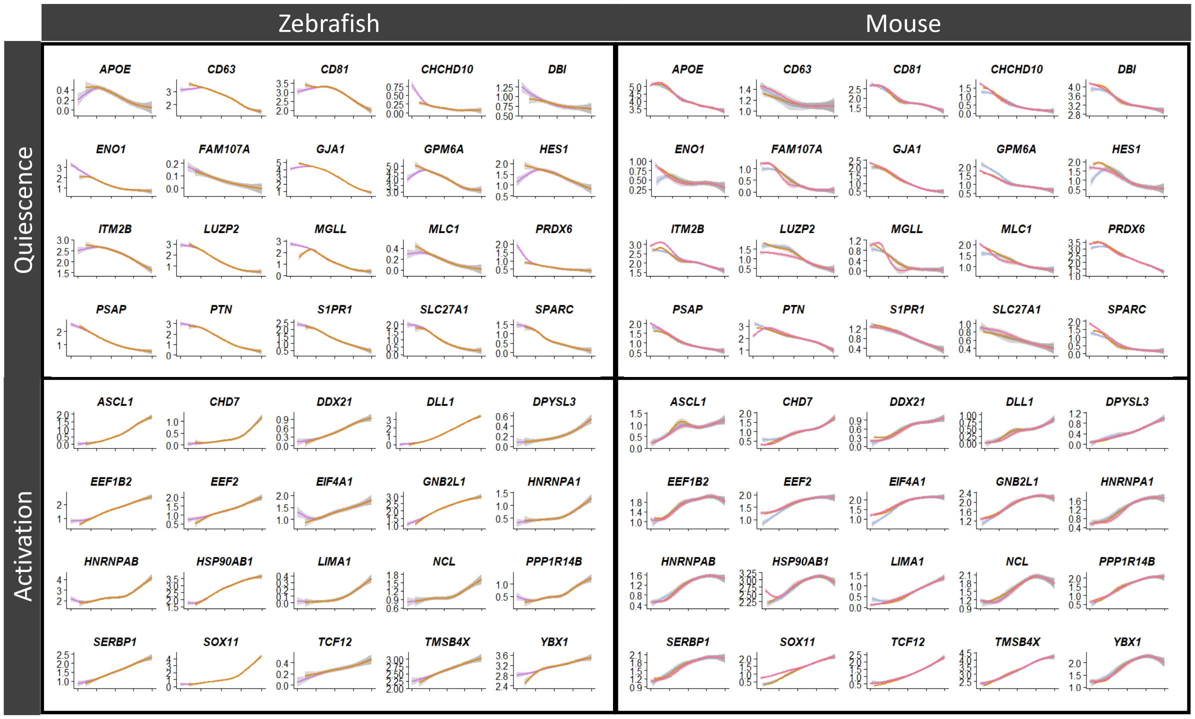


**Fig.S9. Conserved gene expression trajectories in zebrafish and mouse**. Examples of genes with a conserved expression trajectory from deep quiescence to activation (from left to right on the x axis) in the adult zebrafish telencephalon and mouse SEZ. Curve colors match region of origin (see Fig.S8B).

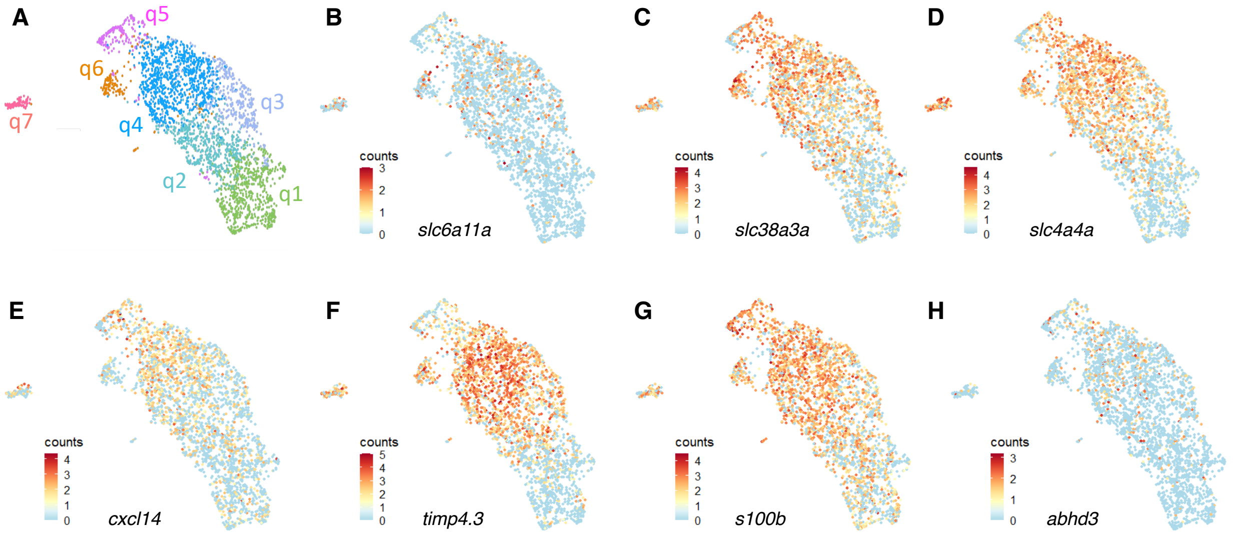


**Fig.S10. Expression of q4-enriched genes. (A)** Annotated UMAP of zebrafish qRG (similar to Fig.2A). **(B-H)** Examples of expression of genes enriched in q4 over q2.

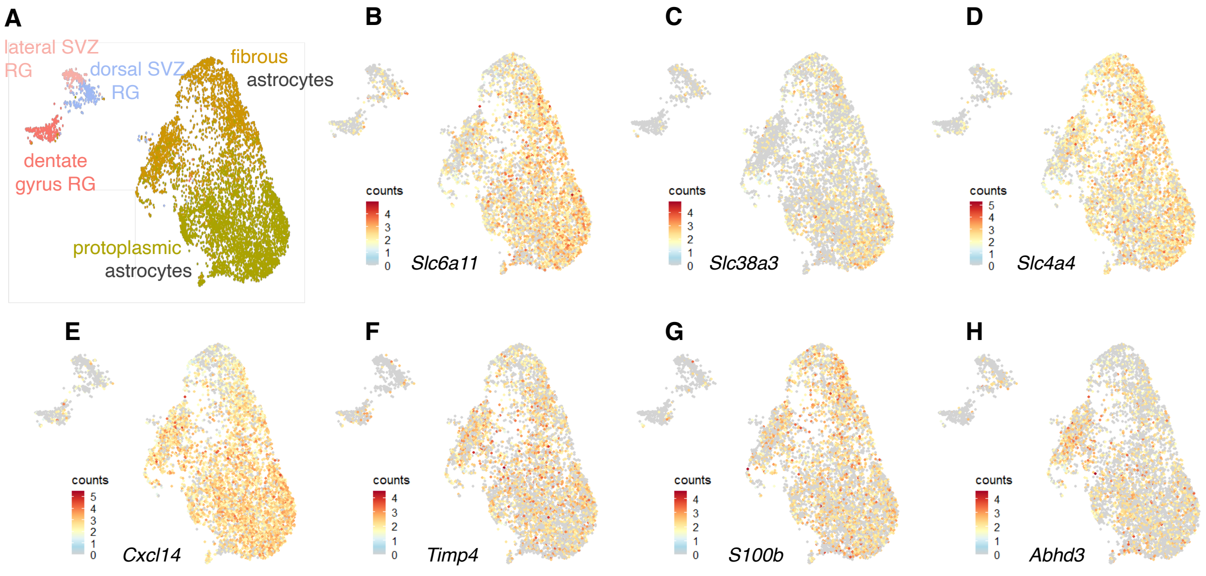


**Fig.S11. Expression of orthologs of zebrafish q4-enriched genes in the mouse telencephalon**. (**A**) Annotated UMAP of telencephalic astroglia from a mouse brain atlas (*49*). (**B-H)** Example of expression of orthologs of genes enriched in zebrafish q4 over q2 plotted in the mouse telencephalon UMAP.

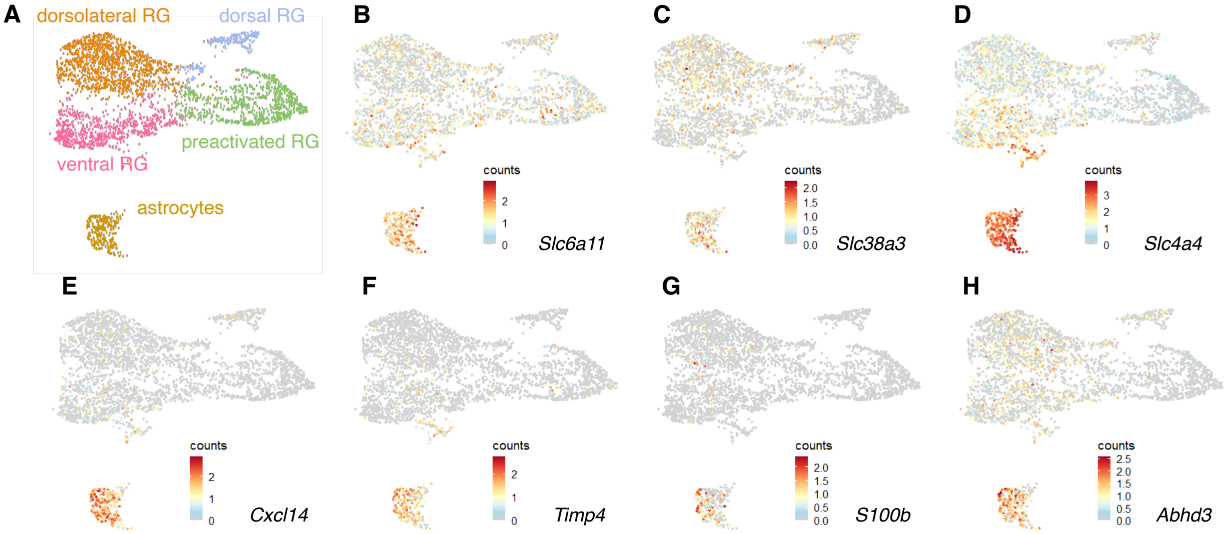


**Fig.S12. Expression of orthologs of zebrafish q4-enriched genes in the mouse SEZ**. **(A)** Annotated UMAP of telencephalic astroglia from mouse SEZ (*10*). **(B-H)** Examples of expression of orthologs of genes enriched in zebrafish q4 over q2 plotted in the mouse SEZ UMAP.

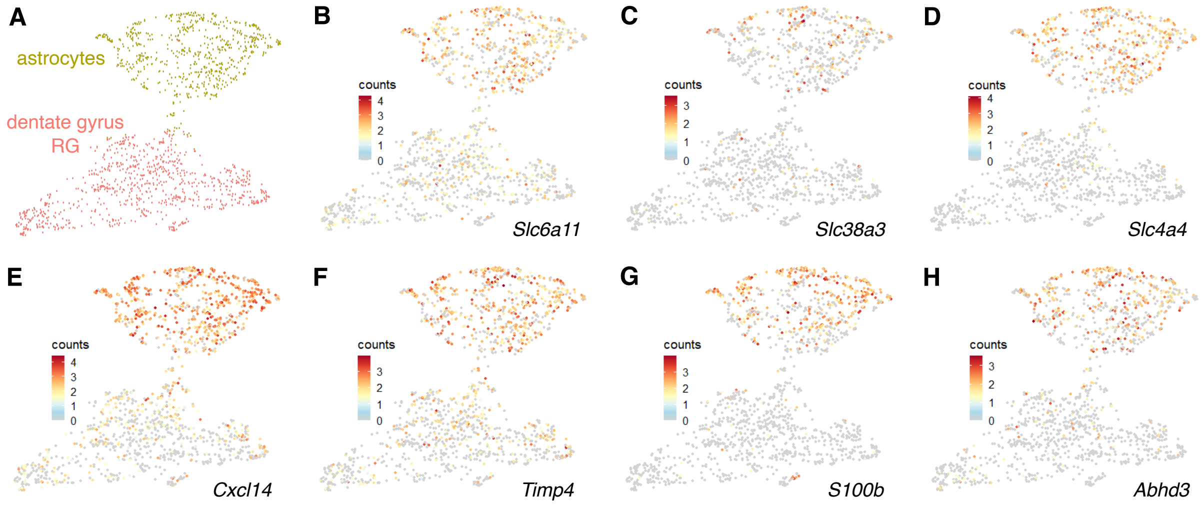


**Fig.S13. Expression of zebrafish orthologs of q4-enriched genes in the mouse hippocampus**. (**A)** Annotated UMAP of astroglia from mouse hippocampus (*32*). (**B-H)** Examples of expression of orthologs of genes enriched in zebrafish q4 over q2 plotted in the mouse hippocampus UMAP.

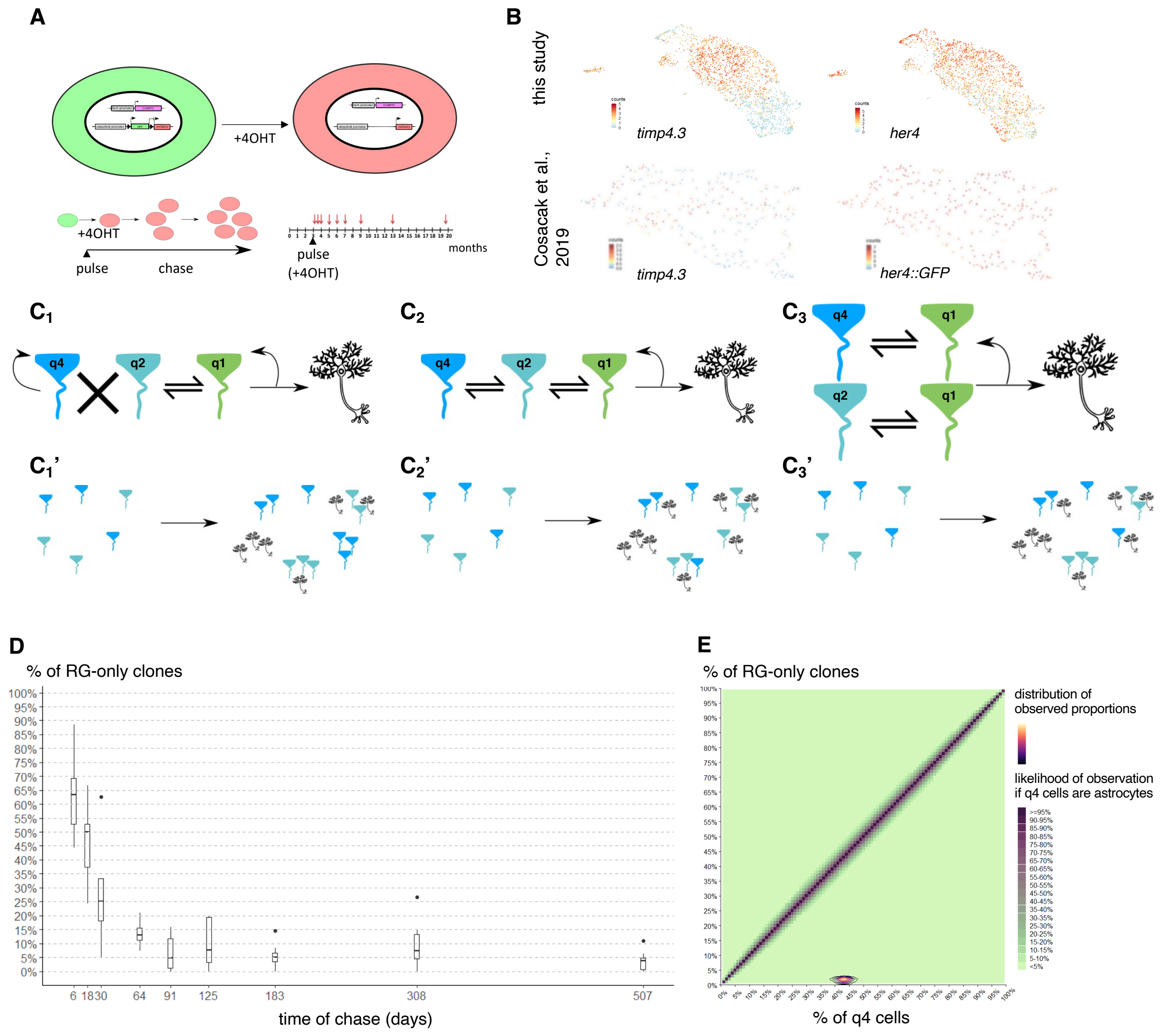


**Fig.S14. Rationale of the analyzed clonal analysis** (76). **(A)** In *Tg(her4:ERT2CreERT2);Tg(ubi:switch)* adult zebrafish, *her4*-positive RG turn on mCherry expression upon clonal tamoxifen induction, and this expression is maintained in their progeny cells, allowing reconstruction of cell lineages after an arbitrarily long chase time. The chronology of analyzed pulse and chase times is depicted. **(B)** Top: this study. *her4* is broadly expressed and enriched in q4 cells suggesting they are targeted by inducible recombination. *timp4.3* expression in our dataset is also shown to highlight q4 cells. Bottom: reanalyzed data from (*25*) showing *her4*-driven *GFP* and *timp4.3*. **(C,C')** Hypotheses on the behavior of q4 RG (C) and outcome after chase (C'). $C_1$-$C_1$': in the first scenario, q4 RG are not neurogenic but are possibly self-replicating like astrocytes. In a clonal analysis labeling both q4 and q2 RG upon induction, the number of clones that have not produced any neurons after chase should be at least equal to the number of q4 RG labeled initially. $C_2$-$C_2$': in the second scenario, q4 RG represent a substate in the quiescence phase of qRG, cells can transit between q4 and q2 and participate in neurogenesis. In a clonal analysis labeling both q4 and q2 cells upon induction, the number of clones that have not given rise to neurons after chase is inferior to the number of q4 cells labeled initially. $C_3$-$C_3$': in the third scenario, q4 and q2 cells are distinct subpopulations but both can activate and participate in neurogenesis. In a clonal analysis labeling both q4 and q2 cells upon induction, the number of clones that have not given rise to neurons after chase is inferior to the number of q4 cells labeled initially. Additionally, clones should be made up of only one type of qRG. (**D)** Time course, per hemisphere, of clones that only contain RG cells, showing a quick decrease followed up by plateauing below 5%. (**E)** Comparison of expected proportions of q4 cells and proportions of non-neurogenic clones. The plot background (green to purple) is colored based on how likely it is to observe a given proportion of clones that do not give rise to astrocytes (y axis) for a given proportion of non-neurogenic cells in the tissue (x axis) as estimated by a bootstrapped Monte Carlo difference of mean test. The density lines (blue to yellow) represent estimations of the actual proportions of non-neurogenic clones and q4 cells and substantially deviate from what would be expected if q4 cells were non-neurogenic.

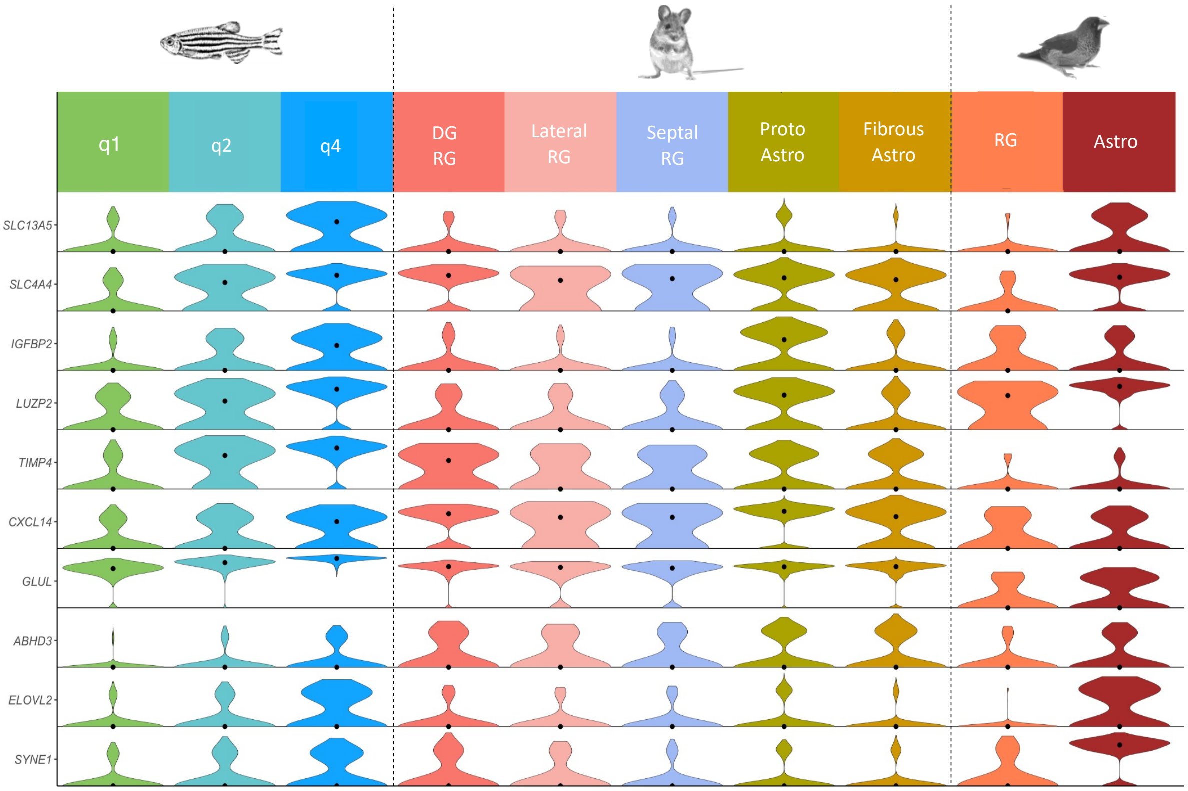


**Fig.S15. Conservation of astrocyte-enriched genes across species**. Violin subset of genes detected as being enriched in q4 over q2 in zebrafish and in astrocytes over RG in mouse, plotted in zebrafish, mouse (*49*) and zebra finch (*55*). Astrocytes from the dentate gyrus were likely not properly separated from RG of the same region, leading to an overestimation of the expression of some transcripts in DG RG here (see Fig.S11).

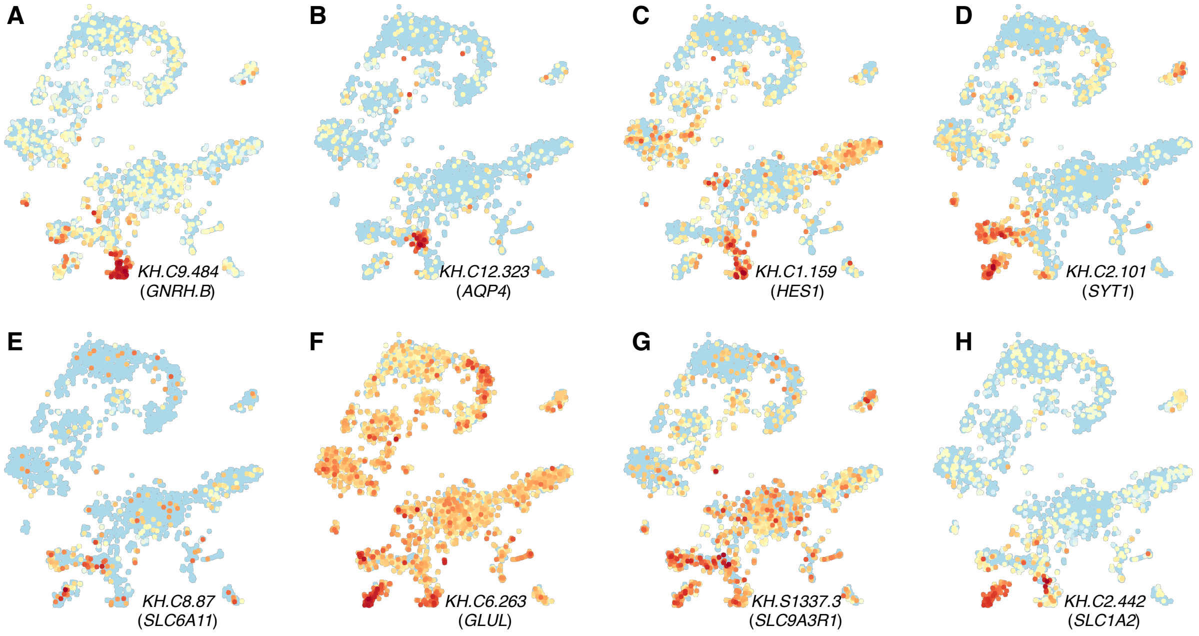


**Fig.S16. Expression of astrocytic genes in Ciona intestinalis swimming larva** (*22*)**, plotted on a scRNAseq UMAP of ciona larval brain cells**. **(A-C)** Expression of genes enriched in ciona ependymoglia (*AQP4* and *HES1* are also markers of mammalian astroglia). **(D)** Expression of a neuronal marker. The cell populations identified with these markers match clusters 5 (ependymoglia) and 3 (differentiated dorsal brain) from the original publication respectively. **(E-H)** Expression of genes belonging to the astrocytic synapomere. Although these genes are enriched in astroglia in vertebrates, they are expressed at higher levels in neurons than in radial ependymoglia in Ciona.

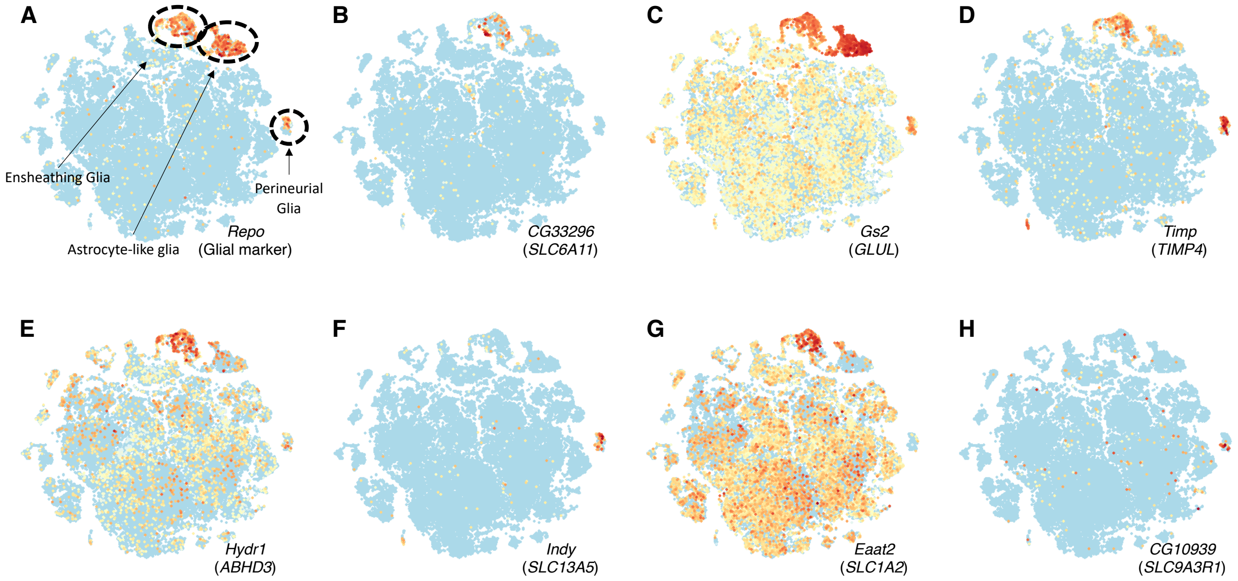


**Fig.S17. Expression of astrocytic genes in Drosophila melanogaster (*16*), plotted on a scRNAseq UMAP of cells isolated from adult Drosophila brains**. *Repo* is used as a general marker of glia, with three major subpopulations highlighted. Expression of orthologs for genes enriched in zebrafish q4 cells and mammalian astrocytes is then depicted, demonstrating their enrichment in ensheathing glia and their distribution across distinct glial populations.

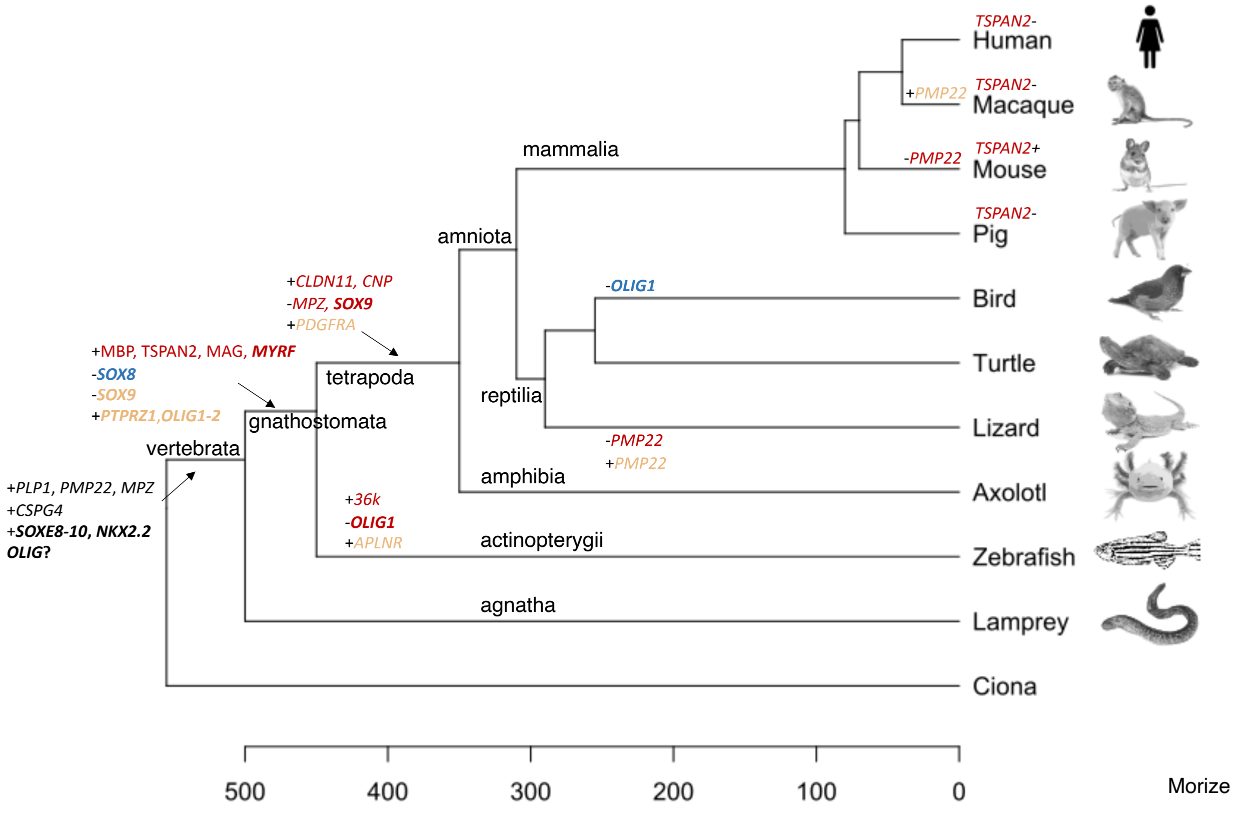


**Fig.S18. Molecular evolution of oligodendrogenesis**. Phylogenetic tree annotated with changes in gene expression inferred from cross-species analysis. Light brown/beige represents changes that happened in OPCs whereas dark brown represents changes in the expression of genes in differentiating/differentiated progeny. Of note this means that some genes represented here can be expressed in daughter cells of OPCs but at different stages of maturation. Changes that happened in both OPCs and their progeny are in blue. Transcription factor-encoding genes are bolded. The expression changes that predate the split between gnathostomes and agnatha is in black without separation between OPCs and their progeny because genes are expressed in the same cell population. We could not distinguish between a loss of *TSPAN2* expression early in mammalian evolution followed by a subsequent gain in rodents or two independent losses in pig and in primates.

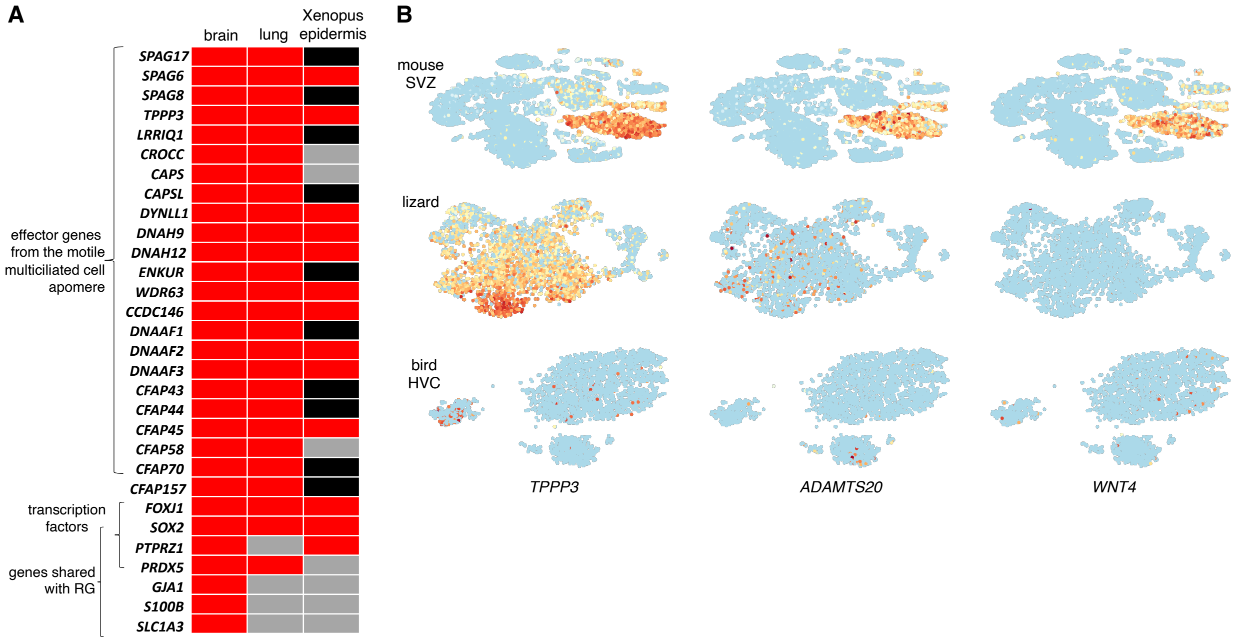


**Fig.S19. Evolution of ependymocytes**. **(A)** Expression of genes related to motile cilia function (top bracket) or shared with astroglia (bottom bracket) in mammalian ependymocytes, human airway ciliated cells (*88*) and xenopus epidermis ciliated cells (*89*). Red squares indicate expression, grey squares indicate lack of expression. In xenopus, black squares indicate that the genes were not detected in any cell cluster in the dataset, which can be either true lack of expression or due to technical artifacts. The top bracket encompasses genes that form a distinctive multiciliated cell apomere that was co-opted several times to produce multiciliated cells with motile cilia in different tissues and species. Despite this convergence, ependymocytes retain exclusive shared expression of several genes with RG, and can be separated from other multiciliated cells based on a limited number of transcription factors, consistent with proposed models of cell-type evolution (*58*). **(B)** De novo expression of niche-modifying factors in mammalian ependymocytes. TPPP3 is used to highlight ependymocytes in three datasets: first a SEZ dataset containing all cell types in mouse, second a dataset of lizard telencephalon containing all astroglial cells and third a bird HVC dataset containing all astroglial cells (distinct astroglial cell types were too hard to distinguish on an embedding of all cells in lizard and bird because these datasets were more diverse and included a smaller proportion of astoglial cells). ADAMTS20 and WNT4 are neurogenesis regulators expressed by ependymocytes in mice but not detected in lizard nor bird ependymocytes.

| | |
|---|---|
| ***SLC1A2*** | Glutamate clearance from the extracellular space |
| ***GLUL*** | Synthesis of glutamine from glutamate, participating in glutamate detoxification |
| *CXCL14* | Related to cytokines |
| *MFGE8* | Inhibition of mTOR, modulation of phagocytosis |
| ***GJA1*** | Gap junction |
| *IGFBP2* | Binding of insulin growth factors |
| *CSPG5* | Proteoglycan |
| *PTN* | Secreted signaling protein |
| *SLC4A4* | Bicarbonate co-transporter involved in pH regulation |
| ***S1PR1*** | Sphingosine receptor |
| ***S100B*** | Calcium binding protein |
| *SEPP1* | Extracellular antioxidant |
| ***SLC6A11*** | GABA uptake transporter modulating the length of Gabaergic transmission |
| *GPM6A* | |
| *TIMP4* | Metallopeptidase inhibitor controlling extracellular matrix composition |
| ***SLC1A3*** | Glutamate transporter involved in termination of glutamate transmission |
| *TSPAN7* | |
| ***APOE*** | Lipidic transporter |
| *PTPLB* | Enzyme involved in the production of very long chain fatty acids |
| ***SLC38A3*** | Sodium-Amino Acid co-transporter, involved in transport of glutamine and neurotransmitters |
| *MGLL* | Converts monoacylglycerides to free fatty acids and glycerol |
| *HEPACAM* | Membrane adhesion protein |
| *ABHD3* | Involved in degradation of medium chain phospholipids |
| *LUZP2* | Leucine zipper transcription factor |
| *SLC9A3R1* | Sodium/proton exchanger with a potential role in ionic and pH control |
| *DIO2* | Involved in local production of active thyroid hormone |
| *ELOVL2* | Involved in elongation of very long chain fatty acids |
| *SYNE1* | Synaptic nuclear envelope protein |
| *SLC13A5* | Sodium-citrate co transporter involved in the regulation of metabolic processes |
| *MTSS1L* | |
| *TMEM176B* | |
| *SLC3A2* | Transporter involved in calcium concentration regulation and transport of amino acids |
| *SLC27A1* | Involved in transcellular lipid transport, in particular of long chain fatty acids |
| *TPI1* | Enzyme involved in glucolysis and gluconeogenesis |
| *ENO1* | Enzyme with glycolytic activity |
| *TTYH3* | Belongs to a family of chlorine ion channels |
| *LAPTM4B* | Involved in lysosomal activity and promotion of autophagy |
| *PHYHIPL* | Predicted to be involved in fatty acid metabolism |
| *PSAP* | Involved in lysosomal activity, principally by promoting catabolism of glycosphingolipids |
| *SPRED1* | Regulator of the MAPK pathway |
| *LRP1* | Involved in lipid homeostasis |
| *EPDR1* | Predicted to be involved in calcium-mediated cell-cell interactions |
| *ASAH1* | Promotes degradation of ceramide into sphingosine and free fatty acids in lysosomes |
| *PLXNB1* | Involved in modulation of cell-cell interactions and control of cell shape |
| *CTSF* | Part of the lysosomal proteolytic system |
| *NDRG3* | |
| ***F3*** | |

**Table S1: Genes with highest confidence of conserved enrichment in q4 over q2 and in astrocytes over RG.**

Gene names are in the left column and known functions pertaining to a potential role in the brain are mentioned in the right column. Genes in bold have been used to label astrocytes in the past with genes in red having been used as distinctive markers between astrocytes and RG in mammalian brains.
